# The mutational landscape of normal human endometrial epithelium

**DOI:** 10.1101/505685

**Authors:** Luiza Moore, Daniel Leongamornlert, Tim H. H. Coorens, Mathijs A. Sanders, Peter Ellis, Kevin Dawson, Francesco Maura, Jyoti Nangalia, Patrick S. Tarpey, Simon F. Brunner, Henry Lee-Six, Raheleh Rahbari, Sarah Moody, Yvette Hooks, Krishnaa Mahbubani, Mercedes Jimenez-Linan, Jan J. Brosens, Christine A. Iacobuzio-Donahue, Inigo Martincorena, Kourosh Saeb-Parsy, Peter J. Campbell, Michael R. Stratton

## Abstract

All normal somatic cells are thought to acquire mutations. However, characterisation of the patterns and consequences of somatic mutation in normal tissues is limited. Uterine endometrium is a dynamic tissue that undergoes cyclical shedding and reconstitution and is lined by a gland-forming epithelium. Whole genome sequencing of normal endometrial glands showed that most are clonal cell populations derived from a recent common ancestor with mutation burdens differing from other normal cell types and manyfold lower than endometrial cancers. Mutational signatures found ubiquitously account for most mutations. Many, in some women potentially all, endometrial glands are colonised by cell clones carrying driver mutations in cancer genes, often with multiple drivers. Total and driver mutation burdens increase with age but are also influenced by other factors including body mass index and parity. Clones with drivers often originate during early decades of life. The somatic mutational landscapes of normal cells differ between cell types and are revealing the procession of neoplastic change leading to cancer.

## Introduction

Acquisition of mutations is a ubiquitous and essential feature of the cells of living organisms. Although there has been comprehensive characterisation of the somatic mutation landscape of human cancer^1-3^, understanding of patterns of somatic mutation in normal cells is limited. In large part this has been due to the challenge of detecting somatic mutations in normal tissues and several strategies have recently been developed to address this including sequencing of *in vitro* derived clonal cell populations from normal tissues^4-8^, sequencing small biopsies containing limited numbers of microscopic clones^9,10^, sequencing microscopically distinguishable structural elements which are clonal units^11,12^, highly error corrected sequencing^13,14^ and sequencing single cells^15,16^. Together, these have begun to reveal differing mutation burdens between different cell types, their patterns of acquisition over time and the signatures of the mutational processes generating them. They have also shown that, in normal tissues, clones of normal cells with “driver” mutations in cancer genes are present. In the glandular epithelium of the colon these are relatively uncommon^12^ but, in the squamous epithelia of the skin^9^ and oesophagus^10^ and other tissues, such as the blood^17-21^, clones carrying drivers can constitute substantial proportions of normal cells present after middle age.

The factors determining differences in mutation landscape between normal cell types are incompletely understood. However, they plausibly include the intrinsic structural and physiological features of each tissue. Endometrium is a uniquely dynamic tissue composed of a stromal cell layer invaginated by a contiguous glandular epithelial sheet covering the luminal surface. It adopts multiple different physiological states during life including premenarche, menstrual cycling, pregnancy, and postmenopause. During reproductive years it undergoes cyclical breakdown, shedding, repair and remodelling in response to oscillating levels of oestrogen and progesterone which entail iterative restoration of the contiguity of the interrupted glandular epithelial sheet that is effected by stem cells within basal glands retained after menstruation^22-25^.

Characterisation of the mutational landscapes of normal tissues is beginning to provide comprehensive understanding of the succession of intermediate neoplastic stages between normal cells and cancers originating from them. There are two major histological classes of endometrial carcinoma^26,27^. Type I, endometrioid carcinoma, is commoner with the main known risk factor being extent of oestrogen exposure, influenced by early menarche, late menopause and body mass index (BMI)^27,28^. Type II, including serous and clear cell carcinomas, occurs in older women with smoking, age and elevated BMI as recognised risk factors^29^. Commonly mutated cancer genes include *PTEN*, *TP53*, *PIK3CA*, *KRAS*, *ARID1A*, *FBXW7* and *PIK3R1^30^* and subsets of endometrial cancer carry large numbers of base substitution and/or small insertion and deletion (indel) mutations due to defective DNA mismatch repair, polymerase epsilon/delta mutations, or large numbers of copy number changes and genome rearrangements^26,31^.

Recent studies using exome and targeted sequencing have revealed the presence of driver mutations in known cancer genes in a high proportion of endometrial glands in endometriosis^11,32,33^, and also in eutopic normal endometrial epithelium^11^. Here, by whole genome sequencing, we have further characterised the mutational landscape of normal endometrial epithelium, explored how it is influenced by age, BMI and parity, estimated the age of driver mutations and the relationship of clonal evolution to glandular architecture.

## Results

### Samples and sequencing

Using laser capture microdissection (LCM) 215 histologically normal endometrial glands were isolated from 18 women aged 19 to 81 years. The samples were from biopsies taken for infertility assessments (6), hysterectomies for benign non-endometrial pathologies (2), residual tissues from transplant organ donors (6) and autopsies after death from non-gynaecological causes (4). DNA from each gland was whole genome sequenced using a library-making protocol modified to accommodate small amounts of input DNA^12^. The mean sequencing coverage was 28-fold and only samples with >15-fold coverage were included in subsequent analyses (n=182, Supplementary Table 1, Supplementary Results 1). Somatic mutations in each gland were determined by comparison with whole genome sequences from pieces of uterus, cervix or Fallopian tube from the same individuals. From each of 18 glands two separate samples were obtained and subjected to independent DNA extraction, library preparation and whole genome sequencing. Using these biological “near-replicates” the mean sensitivity of somatic mutation variant calling was estimated at >86% (range 0.70 – 0.95%) (Methods).

### Clonality of endometrial glands

To assess whether endometrial glands are clonal populations derived from single recent ancestor cells the variant allele fractions (VAFs) of somatic mutations were examined. Most somatic mutations are heterozygous. Heterozygous mutations present in all cells of a population derived from a single ancestor will have VAFs of 0.5 whereas somatic mutations in cell populations derived from multiple ancestors will have lower VAFs or be undetectable by standard mutation calling approaches. 90% (163/182) of microdissected endometrial glands showed distributions of base substitution VAFs with peaks between 0.3 and 0.5 indicating that each consists predominantly of a cell population descended from a single epithelial progenitor stem cell with contamination by other cells potentially including endometrial stromal cells, inflammatory cells and epithelial cells from other glands (Fig. 1, Supplementary Results 1). Similar VAF distributions were observed for small insertions and deletions (indels). Subsequent analyses (see below) have demonstrated that many endometrial glands carry “driver” mutations in known cancer genes. However, endometrial glands exhibited clonality irrespective of the presence of driver mutations with, for example, somatic mutations in all 10 glands from a 19-year-old individual (PD37506) having a median VAF >0.3 but no driver mutations identified (Extended Data Fig. 1a, b). Thus, colonisation of endometrial glands by descendants of single endometrial epithelial stem cells is not contingent on growth selective advantage provided by driver mutations and may occur by a process analogous to genetic drift, as proposed for other tissues^34,35^.

**Figure 1.**
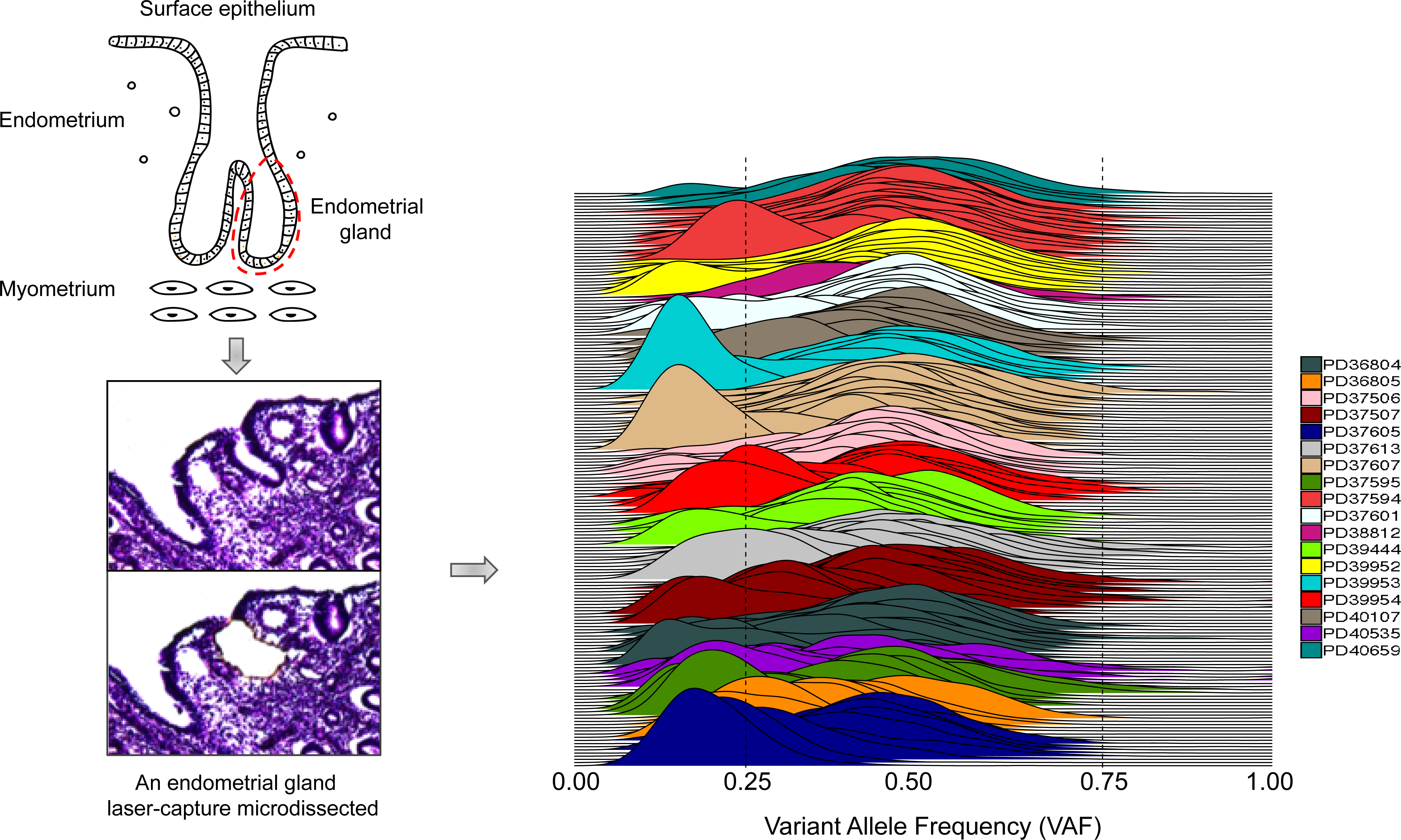
Clonality of normal endometrial glands. Individual normal endometrial glands were laser-capture microdissected and whole genome sequenced. Majority (90%) of the sampled endometrial glands were clonal, i.e. shared most recent common ancestor, with a median variant allele frequency (VAF) between 0.3 and 0.5 for all substitutions.

### Mutation burdens

The somatic mutation burdens in normal endometrial glands from the 18 individuals ranged from 225 to 2890 base substitutions (mean 1324) and 3 to 243 indels (mean 85) (Fig. 2a, b). In large part this variation was attributable to the ages of the individuals with a linear accumulation of ∼28 base substitutions per gland per year during adult life (linear mixed-effect model, SE = 3.1, *P* = 1.061e-07) (Supplementary Results 2). However, the possibilities of lower mutation rates premenarche and postmenopause cannot be excluded. The potential influences of BMI, a known risk factor for endometrial cancer, and the presence of driver mutations on mutation burden were also examined. An additional 20 substitutions were acquired with each unit of BMI (SE = 8, *P* = 2.330e-02). Therefore, the association between elevated BMI and increased endometrial cancer risk may, at least partially, be mediated by this additional mutation burden induced by BMI in normal endometrial epithelial stem cells. Positive driver mutation status conferred an addition of ∼177 substitutions (SE = 45.7, *P* = 1.632e-04). The basis of this correlation is unclear. It is conceivable that an elevated total mutation load increases the chances of including, by chance, a driver. It is also plausible, however, that drivers engender biological changes, for example elevated cell division rates, that result in higher overall mutation loads. There was no obvious correlation between parity and mutation burden.

**Figure 2.**
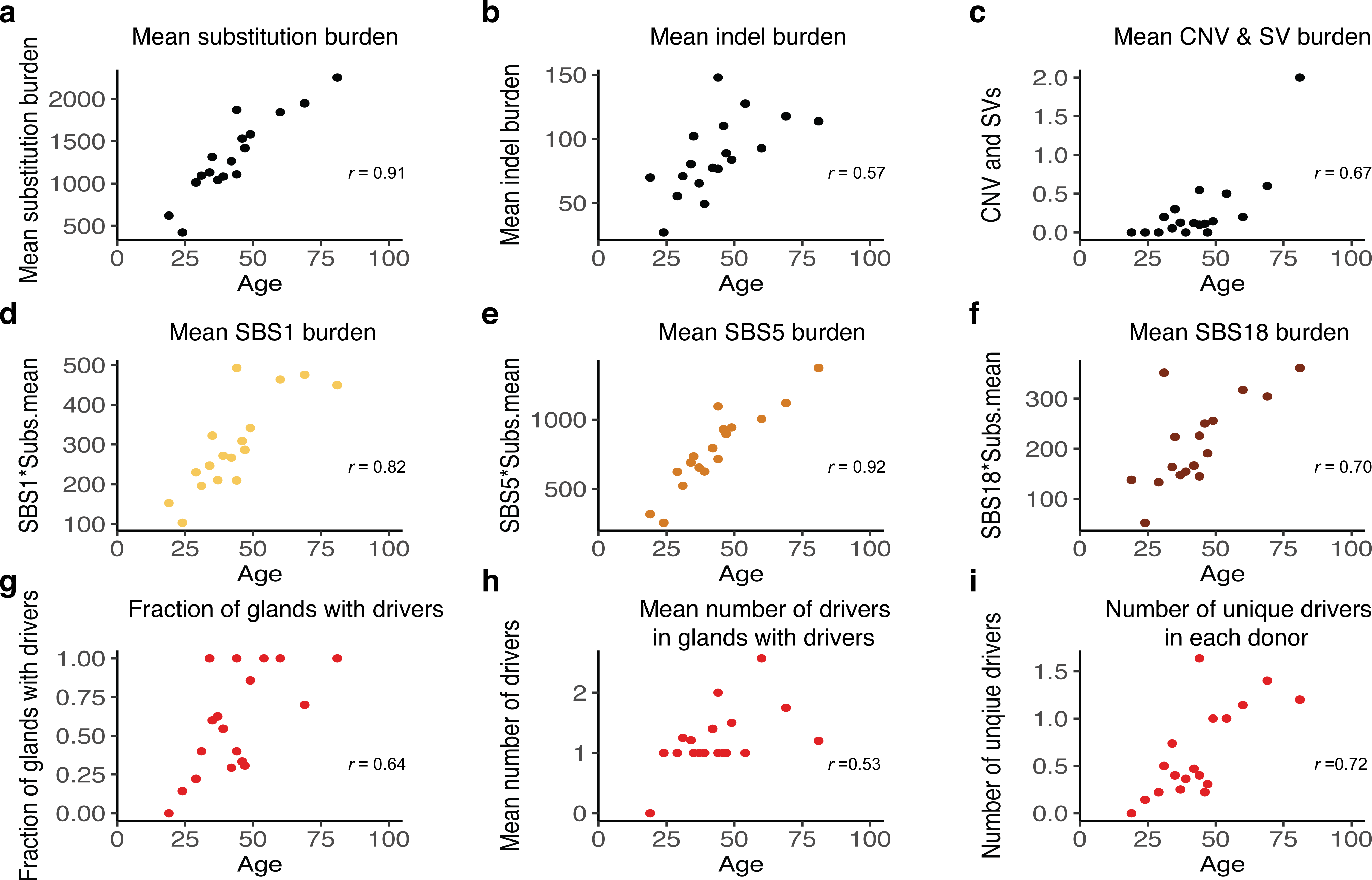
Mutation burden in normal endometrial glands. (**a**) Substitutions accumulate in the endometrium in a relatively linear fashion at an estimated rate of ∼28 substitutions per year (mixed-effect model, *P* = 1.061e-07). A positive correlation between age and accumulation of indels (**b**), copy number and structural variants (**c**) and mutations attributed to mutational signature SBS1 (**d**), SBS5 (**e**) and SBS18(**f**) was also observed. The fraction of glands with driver mutations (**g**), mean number of driver mutations (**h**) and number of unique (different) driver mutations (**i**) all show positive correlation with age.

In addition to endometrial glands, nearby normal endocervical glands were microdissected from one individual (PD37506). There was a ∼2-fold lower somatic mutation burden in endocervical than endometrial glands (Extended Data Fig. 3). This may reflect the absence, in endocervical glands, of the cyclical process of loss and regeneration that occurs in endometrial glands.

**Figure 3.**
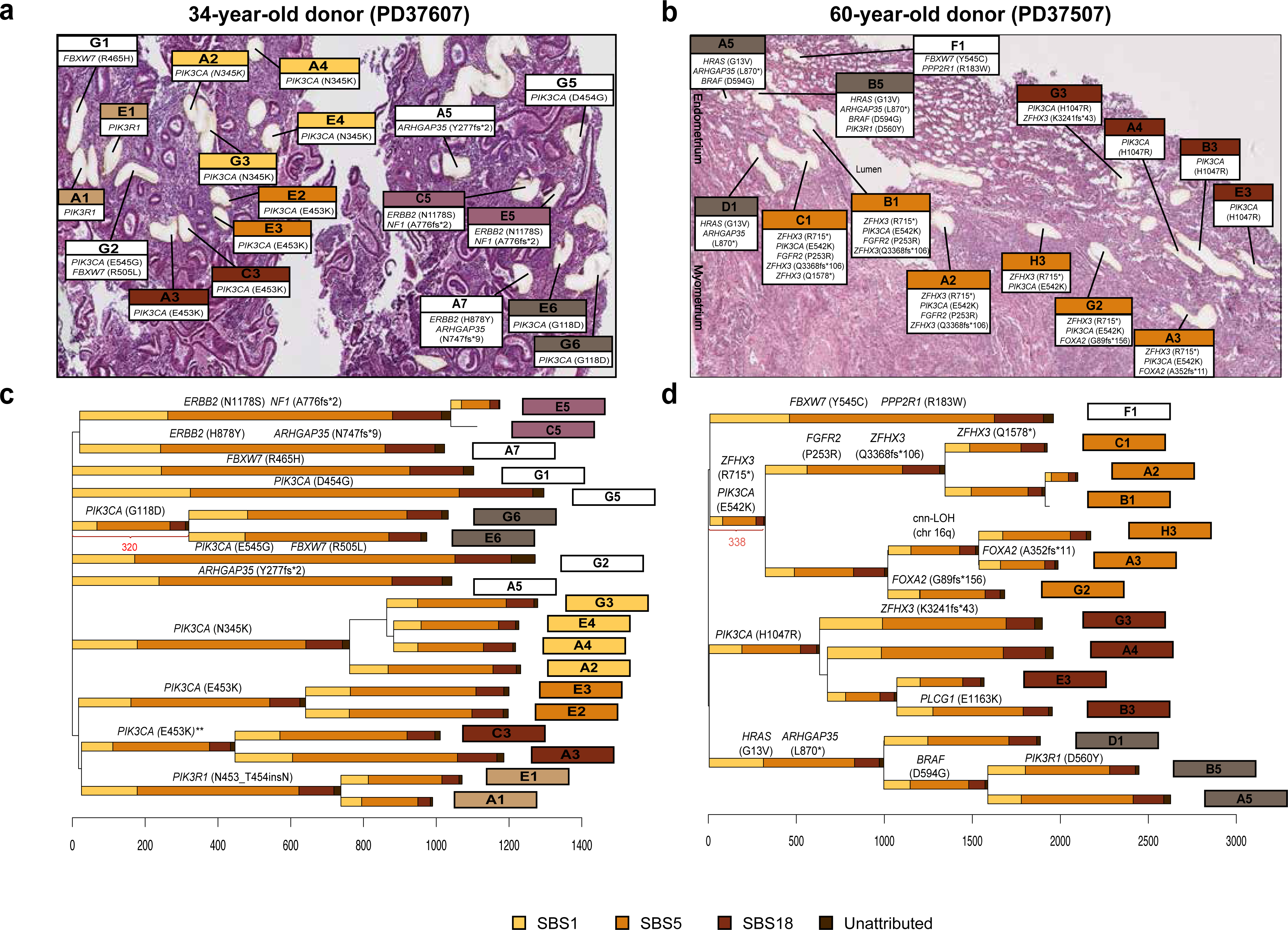
Histology images and phylogenetic trees of normal endometrial glands for two selected individuals with an entirely neoplastic endometrium: 34-year-old (a,b) and 60-year-old (c,d). (**a,b**) Haematoxylin and eosin (H&E) images of individual endometrial glands were taken after laser-capture microdissection (20x magnification). (**c,d**) Phylogenetic trees were reconstructed using single base substitutions; the length of each branch is proportional to the number of variants; a stacked barplot of attributed SBS mutational signatures that contributed to each branch is then superimposed onto every branch; signature extraction was not performed on branches with less than 100 substitutions. The ordering of signatures within each branch is for visualization purposes only as it is not possible to time different signatures within individual branches. Endometrial glands that shared more than 100 variants were considered to belong to the same clade (indicated by the colour of the sample ID label). Labels for glands that did not belong to any clades, are coloured white. The histology images are annotated accordingly. Single base substitution (SBS) signatures are colour-coded (SBS1, SBS5 and SBS18); a small proportion of substitutions across branches were not attributed to reference signatures (‘Unattributed’).

### Mutational signatures

To explore the underlying processes of somatic mutagenesis operative in normal endometrial epithelial cells mutational signatures were analysed. Three previously described single base substitution (SBS) mutational signatures were identified in all endometrial glands (Extended Data Fig. 2): SBS1, predominantly characterised by NCG>NTG mutations and likely due to spontaneous deamination of 5-methylcytosine; SBS5, a relatively featureless, ‘flat’ signature of uncertain cause; SBS18, predominantly characterised by C>A substitutions and possibly due to reactive oxygen species^36^. Overall, the mean signature exposures per gland were 0.22 for SBS1, 0.59 for SBS5 and 0.17 for SBS18; interestingly, glands from one donor with a history of recurrent missed miscarriage (RMM) showed much higher mean SBS18 exposure (0.35) compared to the rest of the cohort. There were approximately 2.7-fold more SBS5 than SBS1 mutations (SD 0.4171666). A positive linear correlation with age for the mutation burden attributable to all three signatures was observed (Fig. 2d, e, f). To ascertain the periods during which different mutational processes operate, phylogenetic trees of endometrial glands were constructed for each individual using somatic mutations (Figs. 3, 4). These revealed that the mutational processes underlying the three signatures are active throughout life. With respect to small indels, composite mutational spectra for each donor were generated and were similar across ages; however, due to the relative sparsity of indels in normal endometrial glands, formal signature extraction was not performed (Extended Data Figure 3).

**Figure 4.**
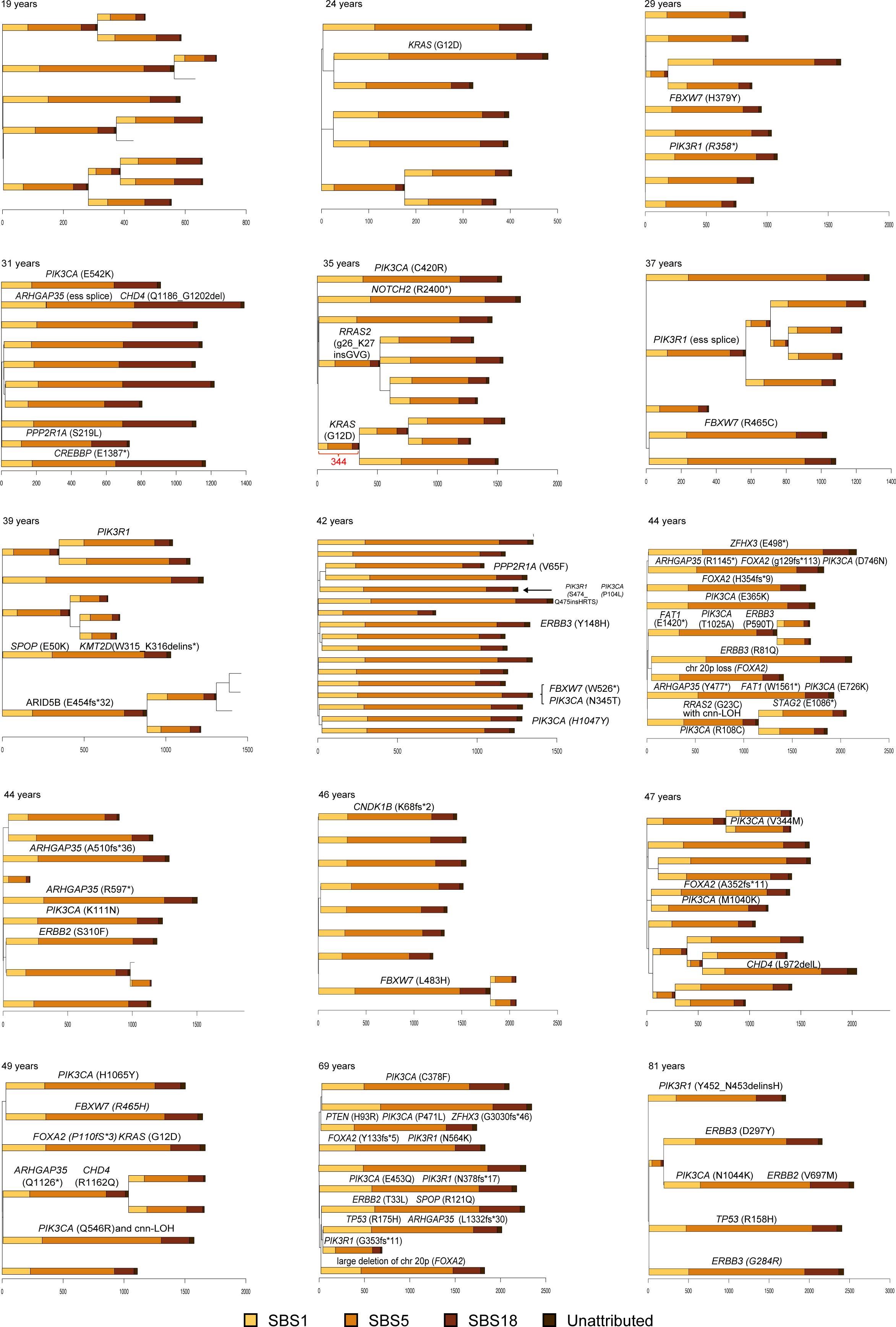
Phylogenetic trees of endometrial glands for all other donors. Phylogenetic trees for the other fifteen donors were reconstructed also using single base substitutions with branch length proportional to the number of variants; the stacked bar plots represent attributed SBS mutational signatures that contributed to each branch. Signature extraction was not performed on branches with less than 100 substitutions. The ordering of signatures within each branch is for visualization purposes only as it is not possible to time different signatures within individual branches.

Somatic copy number changes and structural variants (genome rearrangements) were found in only 27 out of 182 (15%) normal endometrial glands (Fig. 2c, Supplementary Results 3). These included copy number neutral loss of heterozygosity (cnn LOH) in six glands, whole chromosome copy number increases in one and structural variants in eighteen (12 large deletions, six tandem duplications and nine translocations). The majority of glands showed a single change. However, one of two glands carrying a *TP53* mutation (see below) exhibited nine structural variants, indicating that genomic instability caused by defective DNA maintenance occurs in normal cells.

### Driver mutations

To identify genes under positive selection a statistical method based on the observed:expected ratios of non-synonymous:synonymous mutations was used^30^. Eleven genes showed evidence of positive selection in the 182 normal endometrial glands; *PIK3CA*, *PIK3R1*, *ARHGAP35*, *FBXW7*, *ZFHX3*, *FOXA2*, *ERBB2*, *CHD4*, *KRAS*, *SPOP* and *ERBB3* (Supplementary Results 4). All were present in a set of 369 genes previously shown to be under positive selection in human cancer^30^. In addition, four different truncating mutations (and no other mutations) were observed in the progesterone receptor gene (PGR). Although these did not attain standard significance levels the biological role that progesterone plays in normal endometrium as an antagonist of oestrogen driven proliferation raises the possibility that these inactivating mutations confer growth advantage. To comprehensively identify drivers in the 182 endometrial glands, mutations with the characteristics of drivers in each of the 369 genes were sought (Methods).

163 driver mutations were found in normal endometrial glands from 17/18 women (Supplementary Results 5). The youngest carrier was a 24 year old (PD40535) with a *KRAS* G12D mutation in 1/7 glands sampled. 58% (105/182) of endometrial glands carried at least one driver mutation, 19% (35/182) carried at least two and 3% (5/182) carried at least four drivers. Remarkably, in four women, aged 34 (19 glands), 44 (11 glands), 60 (14 glands) and 81 (5 glands), all glands analysed carried driver mutations suggesting that the whole endometrium had been colonised by microneoplastic clones (Figs 3, 4). The fraction of endometrial glands carrying a driver (Fig. 2g), the average number of drivers per gland (Fig. 2h) and the number of different drivers in each individual (corrected for number of glands sampled) (Fig. 2i) all positively correlated with age of the individual. However, there were sufficient outliers from this age correlation to suggest that other factors influence colonisation of the endometrium by driver carrying clones. Using a generalised linear mixed effect model, we found that age has a positive association with the accumulation of driver mutations (coefficient = 0.0336, SE = 0.0131), while parity has a negative association (coefficient = - 0.330, SE = 0.117) (Supplementary Results 6 and 7).

Driver mutations in both recessive (tumour suppressor genes) and dominant cancer genes were found. *PIK3CA* was the most frequently mutated cancer gene, with at least one missense mutation in 61% (11/18) of women and five different mutations found in two women (Fig. 3 and 4, Extended Data Fig. 4). Most truncating driver mutations in recessive cancer genes (including in *ZFHX3*, *ARGHAP35* and *FOXA2* which showed evidence of selection in normal endometrial glands, see above) were heterozygous without evidence of a mutation inactivating the second, wild type allele. Therefore, haploinsufficiency of these genes appears sufficient to confer growth advantage in normal cells. Nevertheless, further inactivating mutations, including copy number neutral LOH of the wild type allele and truncating mutations, in the same genes in other glands indicate that additional advantage is conferred by complete abolition of their activity (notably for *ZFHX3* in the 60 year old, Figure 3 and Exended Data Fig. 5). Driver mutations were found in genes encoding growth factor receptors (*ERBB2*, *ERBB3, FGFR2*), components of signal transduction pathways (*HRAS*, *KRAS*, *BRAF*, *PIK3CA*, *PIK3R1*, *ARHGAP35*, *RRAS2*, *NF1*, *PP2R1A*, *PTEN*), pathways mediating steroid hormone responses (*ZFHX3*, *FOXA2*, *ARHGAP35*), pathways mediating WNT signalling (*FBXW7*) and proteins involved in chromatin function (*KMT2D*, *ARID5B*). Many different combinations of mutated cancer genes were found in individual glands.

**Figure 5.**
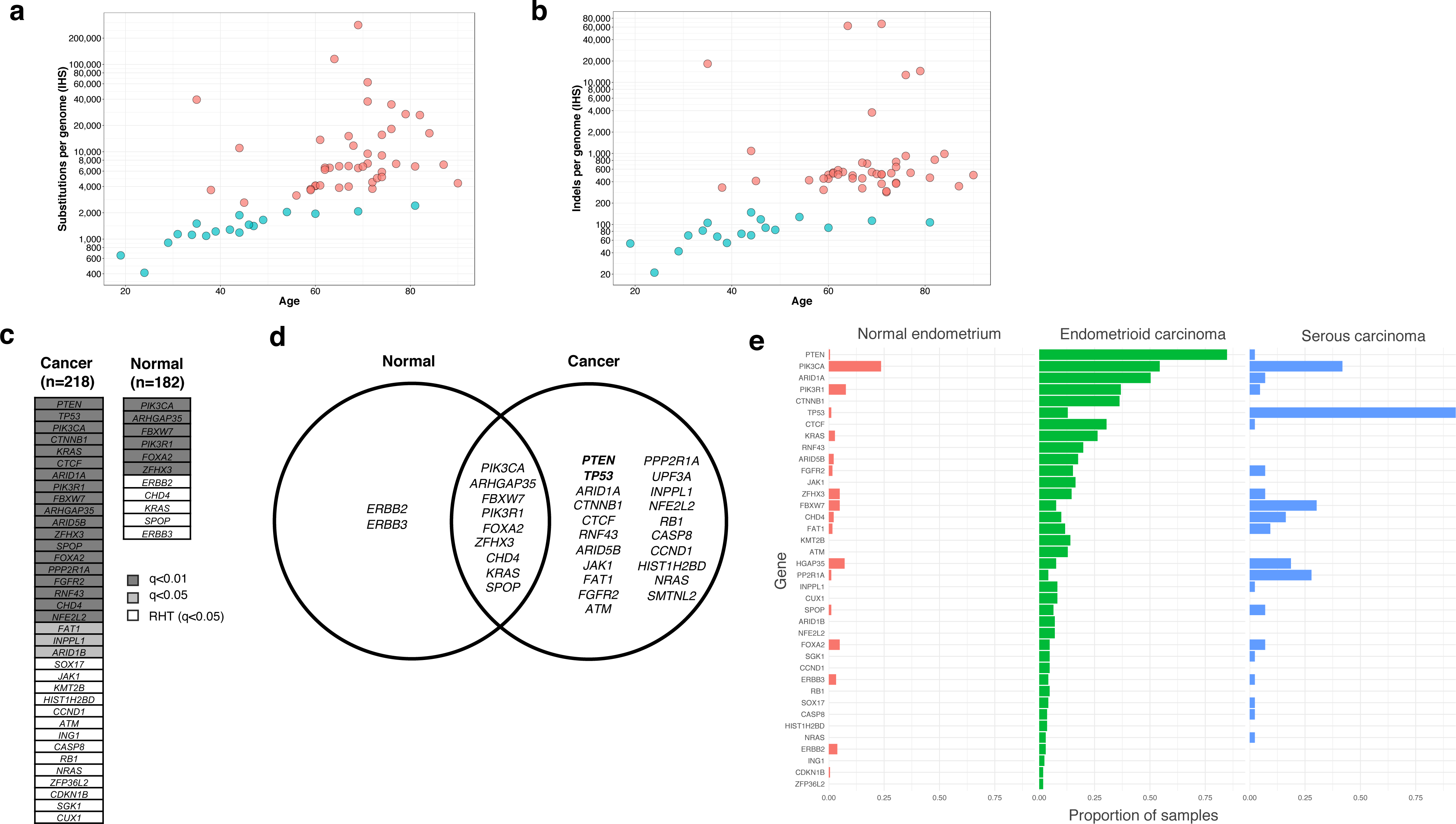
Comparison between normal endometrial epithelium and endometrial cancer. (**a**,**b**) Normal endometrial glands show lower total mutation burden in comparison to endometrial cancer. (**d,e**) Genes that are under significant positive selection (dn/ds>1) in normal endometrial epithelium and endometrial cancer. RHT, restricted hypothesis testing of known cancer genes. *ERBB2* and *ERBB3* are under selection in normal endometrial epithelium, but are not in endometrial cancer. (**f**) Identified driver mutations and their distribution in normal endometrial glands and the two major types of endometrial cancer (endometroid and serous carcinomas).

Driver mutations were placed on the phylogenetic trees of somatic mutations constructed for each individual and, by assuming a constant somatic mutation rate during life, the time of occurrence of a subset was estimated (Methods). Some driver mutations occurred early in life. These included a *KRAS* G12D mutation in three glands from a 35 year old and a *PIK3CA* mutation in two glands from a 34 year old, which are both likely to have arisen during the first decade. A pair of drivers in *ZFHX3* and *PIK3CA*, co-occurring in six glands from a 60 year old, was also acquired during the first decade indicating that driver associated clonal evolution begins early in life. There was evidence, however, for continuing acquisition and clonal expansion of driver mutations into the third and fourth decades and further accumulation beyond this period is not excluded.

### Phylogeography of mutations within the endometrium

Phylogenetically closely related glands were often in close physical proximity within the endometrium (Fig. 3). In phylogenetic clusters for which the mutation catalogues were almost identical, this may simply reflect multiple sampling of a single tortuous gland weaving in and out of the plane of section, rather than distinct glands with their own stem cell populations (e.g. glands C5 and E5, Figs. 3a, c). For other phylogenetic clusters, the different branches within the clade have diverged substantially, sometimes acquiring different driver mutations, and therefore are likely derived from different stem cell populations. In such instances phylogenetically related glands can range over distances of multiple millimetres suggesting that their clonal evolution has entailed capture and colonisation of extensive zones of endometrium (e.g. glands C1, A2, B1, H2, A3, B3, Figs. 3b, d). Conversely, many glands in close physical proximity are phylogenetically distant (e.g. glands E1 and G2, Figs 3a, c), indicating that the cell populations have remained isolated from each other.

### Normal endometrium compared to other cells

Endometrial cells exhibit lower mutation rates than normal skin epidermal^9^, colorectal^4,12^, small intestinal^4,12^ and liver cells^4^, similar burdens to oesophageal cells^10^ and higher rates than skeletal muscle cells^7^ (Extended Data Fig. 7). Of the mutational signatures found in endometrial cells, SBS1 and SBS5 are found in all other cell types^37^. However, the SBS1 mutation rate is higher in colorectal and small intestinal epithelial cells whereas the SBS5 mutation rate is higher in liver cells^4^. SBS18 has also been found ubiquitously in colonic crypts^12^.

The prevalence of driver mutations is substantially different in different normal cell types. In colon, like the endometrium a tissue with glandular architecture, only ∼1% crypts (glands) in 60 year old individuals carry a driver mutation^12^ compared to the much higher fractions (up to 100%) in the endometrial glands of 60 year old women. The biological basis of this difference is unclear but is unlikely to be the difference in total mutation burden, which is lower in the endometrium than the colon.

Endometrial cancers exhibit higher mutation loads than normal endometrial cells, for base substitutions (∼5-fold, medians of 1346 and 7330 substitutions observed in normal endometrium and endometrial cancer respectively (Mann-Whitney test, *P* = 7.629e-06) (Fig. 5a)) and indels (Fig. 5b) and these differences also pertain to normal endometrial cells with driver mutations. In most endometrial cancers these differences are attributable to higher mutation burdens of the ubiquitous base substitution and indel mutational signatures. In addition, however, the very high mutation loads of the subsets of endometrial cancers with DNA mismatch repair deficiency and polymerase epsilon/delta mutations were not seen in normal endometrial cells. Differences between endometrial cancers and normal cells were even more marked for structural variants and copy number changes (median number zero in normal endometrial cells and ∼23 in endometrial cancers^38^) and this again pertained to normal endometrial cells with drivers.

There were also differences in the repertoire of cancer genes in which driver mutations were found (Fig 5 d,e,f, Supplementary Results 4 and 8). Notably, mutations in *PTEN*, *CTCF*, *CTNNB1* and *ARID1A* in endometrioid and in *TP53* in serous endometrial cancer accounted for higher proportions of driver mutations than in normal endometrial cells. It is possible that *PTEN*, *ARID1A*, *TP53* and *CTCF* require biallelic mutation to confer growth advantage and this may account for their lower prevalence in normal cells. However, heterozygous mutations in *PTEN* and *TP53* were found, albeit rarely and restricted to the two oldest individuals studied (69 and 81-year old), and this explanation would not account for the relative deficit of *CTNNB1* mutations. Overall, the results suggest that driver mutations in some cancer genes may be relatively effective at enabling stem cell colonisation of normal tissues but confer limited risk of conversion to invasive cancers. Conversely, others may require biallelic mutation and/or confer limited advantage in colonising normal tissues but are relatively effective at conversion to malignancy.

## Discussion

This study of normal endometrial epithelium, together with recent studies of other normal cell types^4,5,9-12,17,18^, is revealing the landscape of somatic mutations in normal human cells. The landscape is characterised by different somatic mutation rates in different cell types that, for the most part, are generated by a limited repertoire of ubiquitous mutational processes generating base substitutions, small indels, genome rearrangements and whole chromosome copy number changes. These processes exhibit more or less constant mutation rates during the course of a lifetime resulting in essentially linear accumulation of mutations with age. However, the influences of BMI and the presence of driver mutations on mutation burden in endometrial epithelium indicate that additional factors can modulate their mutation rates. The reasons for the different mutation rates of ubiquitous signatures in different tissues are unclear. For SBS1, which is likely due to deamination of 5-methylcytosine, the differences may be related to the number of mitoses a cell has experienced. Additional mutational signatures which are present only in some cells, only in some cell types and/or are intermittent also operate in normal cells, supplementing the mutation load contributed by ubiquitous signatures. The latter include exposures such as ultraviolet light in skin^9^, APOBEC mutagenesis in occasional colon crypts and other signatures of unknown cause in normal colon epithelium^12^.

A small subset of mutations generated by these mutational processes have the properties of driver mutations. The total somatic mutation rate is lower in endometrial than colonic epithelial stem cells and thus the rate of generation of driver mutations is also likely to be lower. However, numerous cell clones with different driver mutations, some carrying multiple drivers, colonise much of, and in some cases potentially all of, the normal endometrial epithelium in most women. This is in marked contrast to the colon where just 1% of normal crypts in middle-aged individuals carry a driver^12^. This dramatic difference may be due to intrinsic differences in physiology between endometrium and colon. In the endometrium, the cyclical process of tissue breakdown, shedding and remodelling iteratively opens up denuded terrains for pioneering clones of endometrial epithelial cells with drivers to preferentially colonise compared to wild type cells. By contrast, in the colon the selective advantage of a clone with a driver is usually confined to the small siloed population of a single crypt, with only occasional opportunities for further expansion. Thus, the endometrium in some respects resembles more the squamous epithelia of skin and oesophagus in which cell clones derived from basal cells directly compete against each other for occupancy of the squamous sheet and in which substantial proportions of such sheets become colonised over a lifetime by normal cell clones carrying driver mutations^9,39^. Although this rampant colonisation by driver clones in endometrium progresses with age, it is already well advanced in some young women, and parity apparently has an inhibitory effect on it, indicating that multiple factors influence its progression. More extensive studies of the mutational landscape in normal endometrium are required to better assess how pregnancy, the premenarchical and postmenopausal states, hormonal contraceptive use and hormone replacement therapies influence it and also the potential impact it has on pregnancy and fertility.

The burdens of all mutation classes are lower in normal endometrial cells, including those with drivers, than in endometrial cancers. However, these differences are most marked for structural variants/copy number changes and for the extreme base substitution/indel hypermutator phenotypes due to DNA mismatch repair deficiency and polymerase delta/epsilon mutations which were not found in normal endometrium. The results therefore suggest that in endometrial epithelium, and in other tissues thus far studied including colon, oesophagus and skin, normal mutation rates are sufficient to generate large numbers of microneoplastic clones with driver mutations behaving as normal cells, but that acquisition of an elevated mutation rate and burden is associated with further evolution to invasive cancer. Given that the endometrial epithelium is extensively colonised by clones of normal cells with driver mutations in middle-aged and older women and that the lifetime risk of endometrial cancer is only 3%^40^, this conversion from microneoplasm to symptomatic malignancy appears to be extremely rare. Driver mutations in normal endometrium often appear to arise and initiate clonal expansion early in life. It is therefore plausible that some neoplastic clones ultimately manifesting as cancer were initiated during childhood, although the fraction to which this might apply is unclear.

This study has added endometrial epithelial cells to the set of normal cell types in which the landscape of somatic mutations has been characterised. However, most normal tissues have not been investigated in this way. The outcomes of the current studies showing differences in mutation burdens, mutational signatures and prevalence of driver mutations mandates a systematic characterisation of the somatic mutation landscape in all normal human cell types.

## ACKNOWLEDGEMENTS

This work was supported by the Wellcome Trust. LM is a recipient of a CRUK Clinical PhD fellowship (C20/A20917) and Pathological Society of Great Britain and Ireland Trainee Small Grant (Grant Reference No 1175). SFB was supported by the Swiss National Science Foundation (P2SKP3-171753 and P400PB-180790). MAS is supported by a Rubicon fellowship from NWO (019.153LW.038).

We thank Laura O’Neil, Calli Latimer and Paul Scott for technical support; Feran Nadeu and Jingwei Wang for their advice on mutational signature extraction; Thomas J Mitchell, Nicola Roberts and Andrew R.J. Lawson for their assistance with data analysis. We are also grateful to the Cambridge Biorepository for Translational Medicine for the provision of samples from deceased transplant organ donors.

## AUTHOR CONTRIBUTIONS

MRS and LM designed the study and wrote the manuscript with contributions from all authors. KSP, CAID, JJB, KM, MJL and LM obtained samples. PE and LM devised the protocol for laser-capture microscopy, DNA extraction and sequencing of endometrial glands. LM prepared sections, reviewed histology, micro-dissected and lysed endometrial glands. YH assisted with tissue processing and section preparation. LM performed data curation and analysis with the help from DL, THHC, MAS, KD, JN, PST, SFB, HLS, and RR. THHC reconstructed phylogenetic trees. MAS devised filters for substitutions and structural variants. DL, FM and SM assisted with signature analyses. IM assisted with statistical and dnds analyses. PJC oversaw statistical analyses and performed analysis of structural variants. MRS supervised the study.

## EXTENDED FIGURE LEGENDS

**Extended Data Figure 1.**
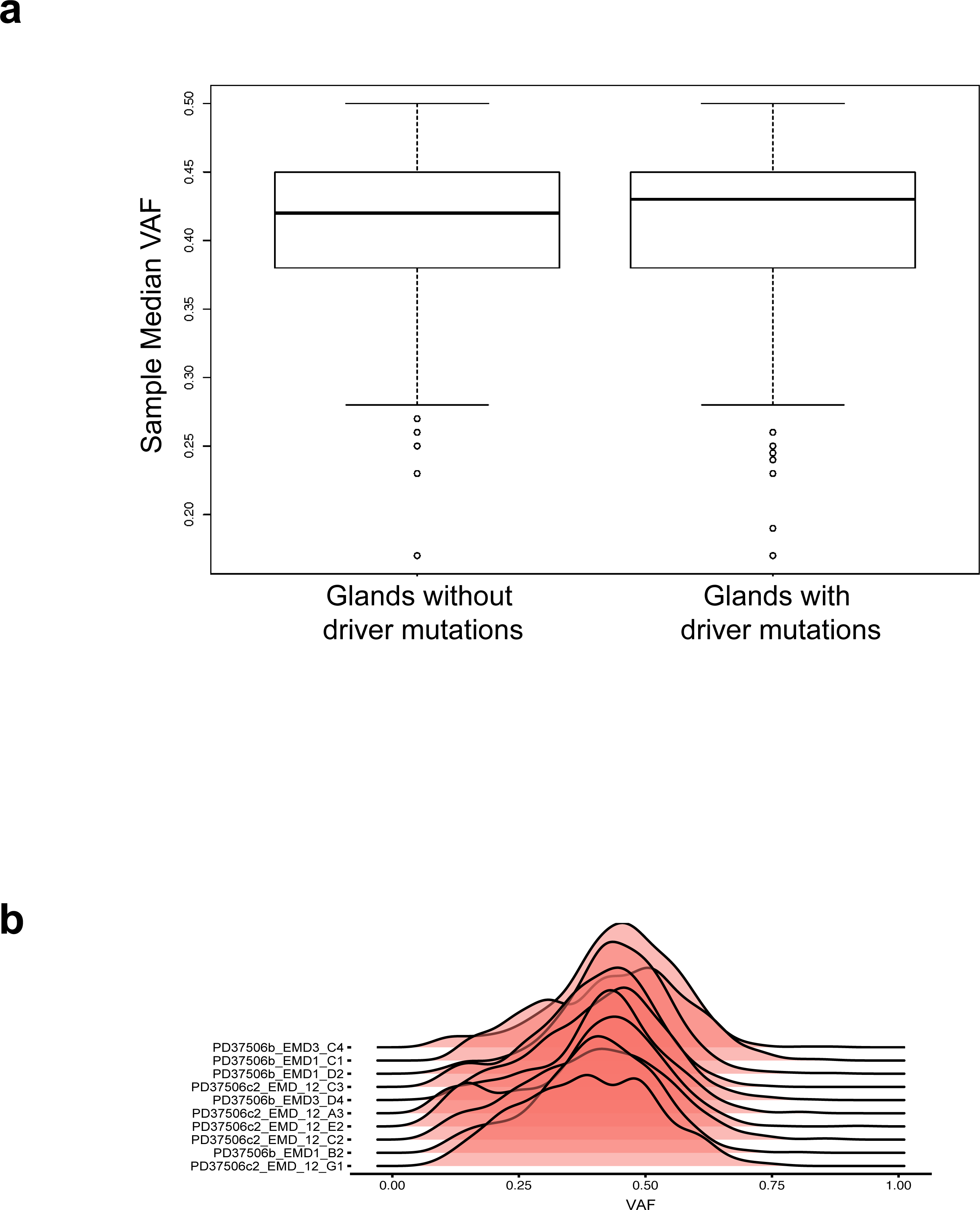
Clonality of endometrial glands and driver mutations. (**a**) The majority of sampled normal endometrial glands were clonal with a median variant allele frequency (VAF) of 0.3 or above. The observed monoclonality of the glands was independent of the driver status (Mann-Whitney two-sided test, *P* = 0.1999). (**b**) All glands from the 19-year-old donor (PD37506) were clonal with a median VAF >=0.3, but there were no detectable driver mutations.

**Extended Data Figure 2.**
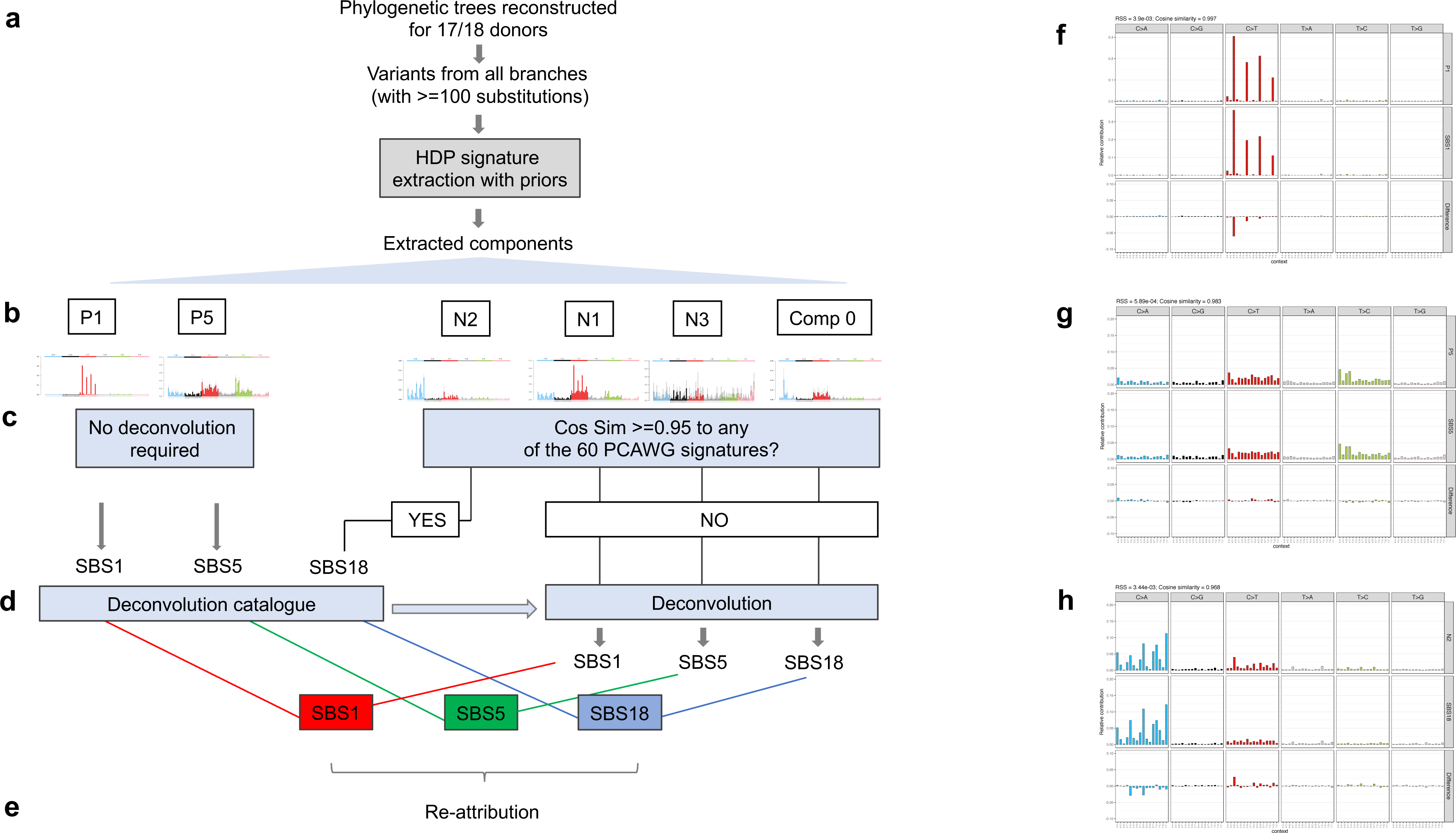
Single Base Substitution (SBS) mutational signatures in normal endometrial glands. (**a**) Final catalogue of single base substitutions were used to re-construct phylogenetic trees for 17 donors (due to the low number of high depth samples, genomes from donor PD38812 were not included in this analysis). SBS signatures were extracted on a per branch basis using a Hierarchical Dirichlet Process (HDP) with a set of 19 reference signatures that were identified in endometrial cancer (‘priors’) by the Mutational Signatures working group of the Pan Cancer Analysis of Whole Genomes (PCAWG). **(b)** HDP extracted components included the following: ‘priors’/reference SBS signatures (P1 = SBS1 and P5 = SBS5); ‘new’ components that did not match any of the provided 19 reference signatures/priors (N1, N2 and N3) and ‘Component 0’ (Comp 0). **(c)** As P1 and P5 showed high cosine similarity (>0.95) to SBS1 and SBS5 signatures respectively (**f,g**), no further deconvolution of these components was required. All other extracted components were compared to the full set of 60 reference signatures. If a component had a cosine similarity of >0.95 to any of the reference signatures (N2 = SBS18, **h)**, no further deconvolution was required (**d**). If a component did not show high cosine similarity to any of the reference signatures, deconvolution was performed using a ‘deconvolution’ catalogue comprising all of the extracted signatures (SBS1, SBS5 and SBS18). **(e)** Final exposures were derived and signatures re-attributed to the individual branches.

**Extended Data Figure 3.**
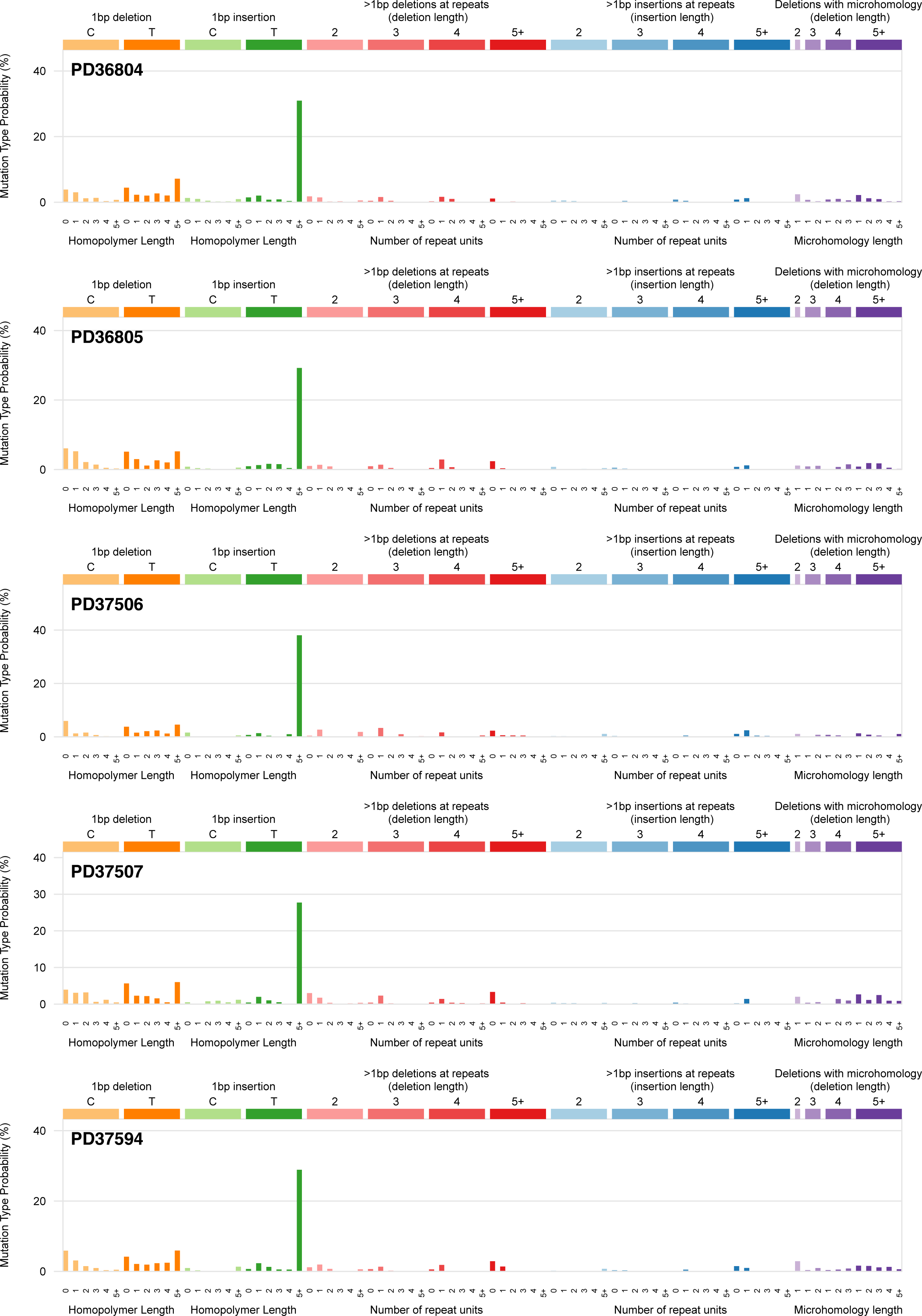
Composite mutational spectra of small insertions and deletions (indels) for each donor. Indels were classified and composite mutational spectra for each individual were generated; due to the relative sparsity of indels detected, no formal signature extraction was performed.

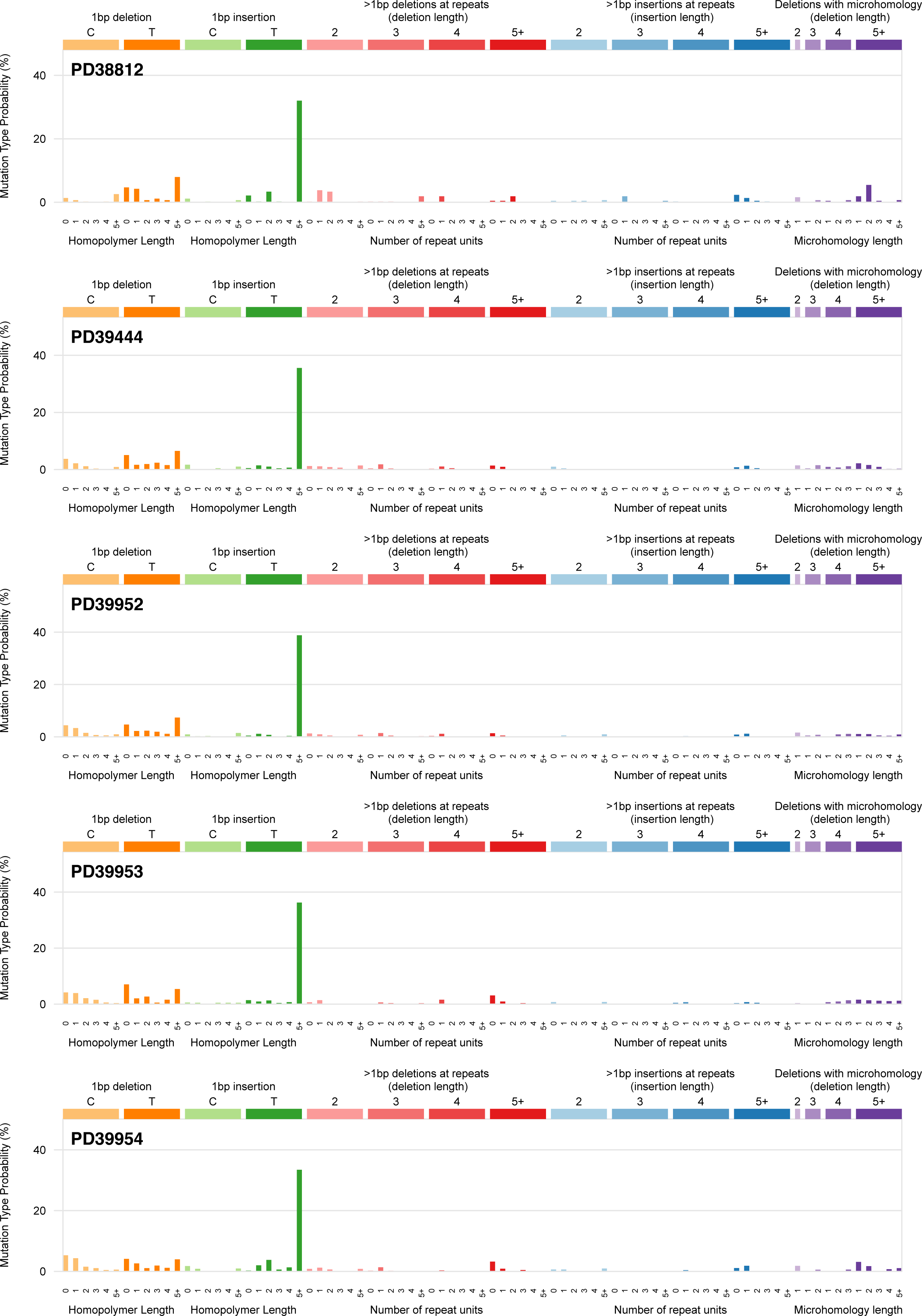

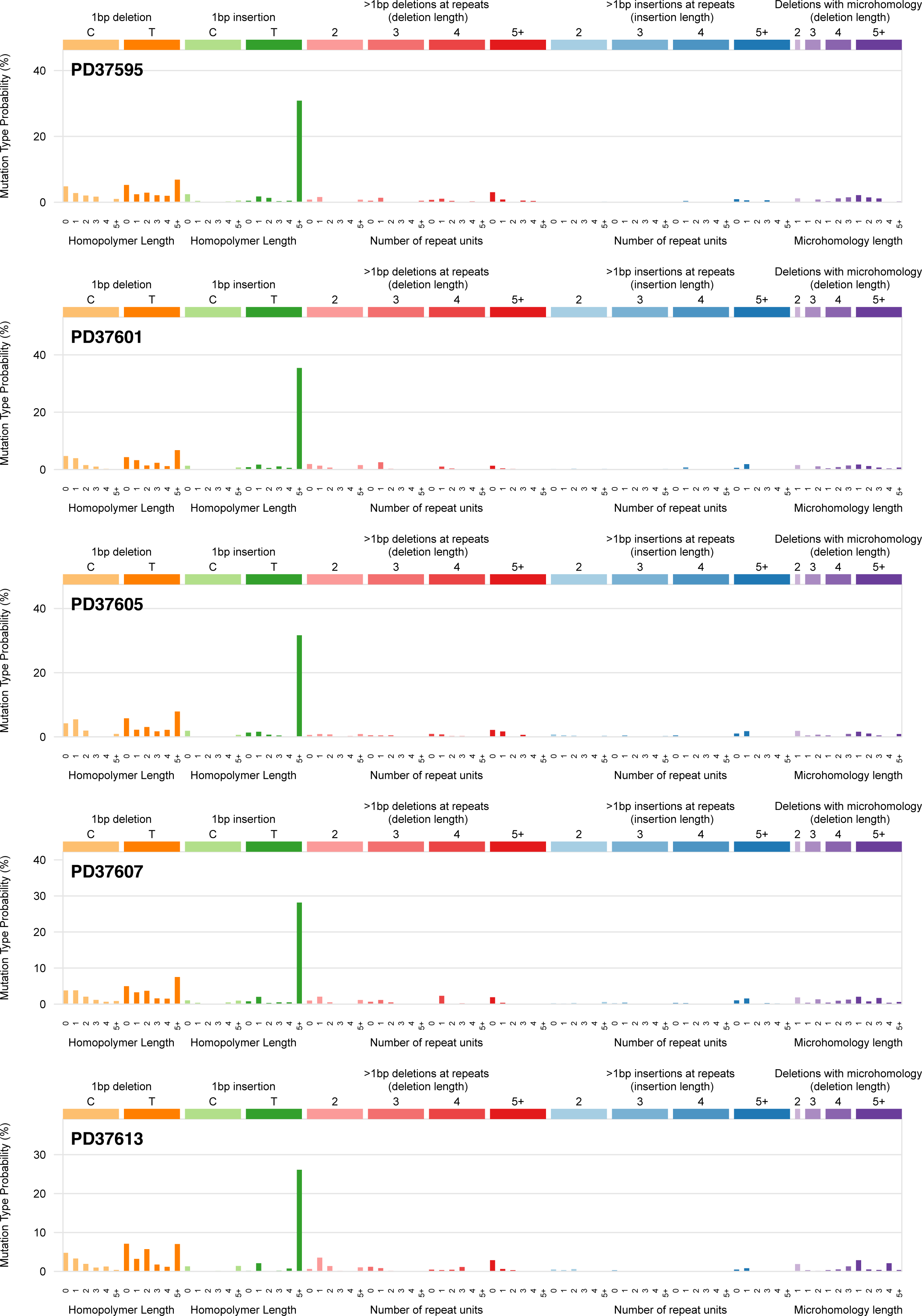

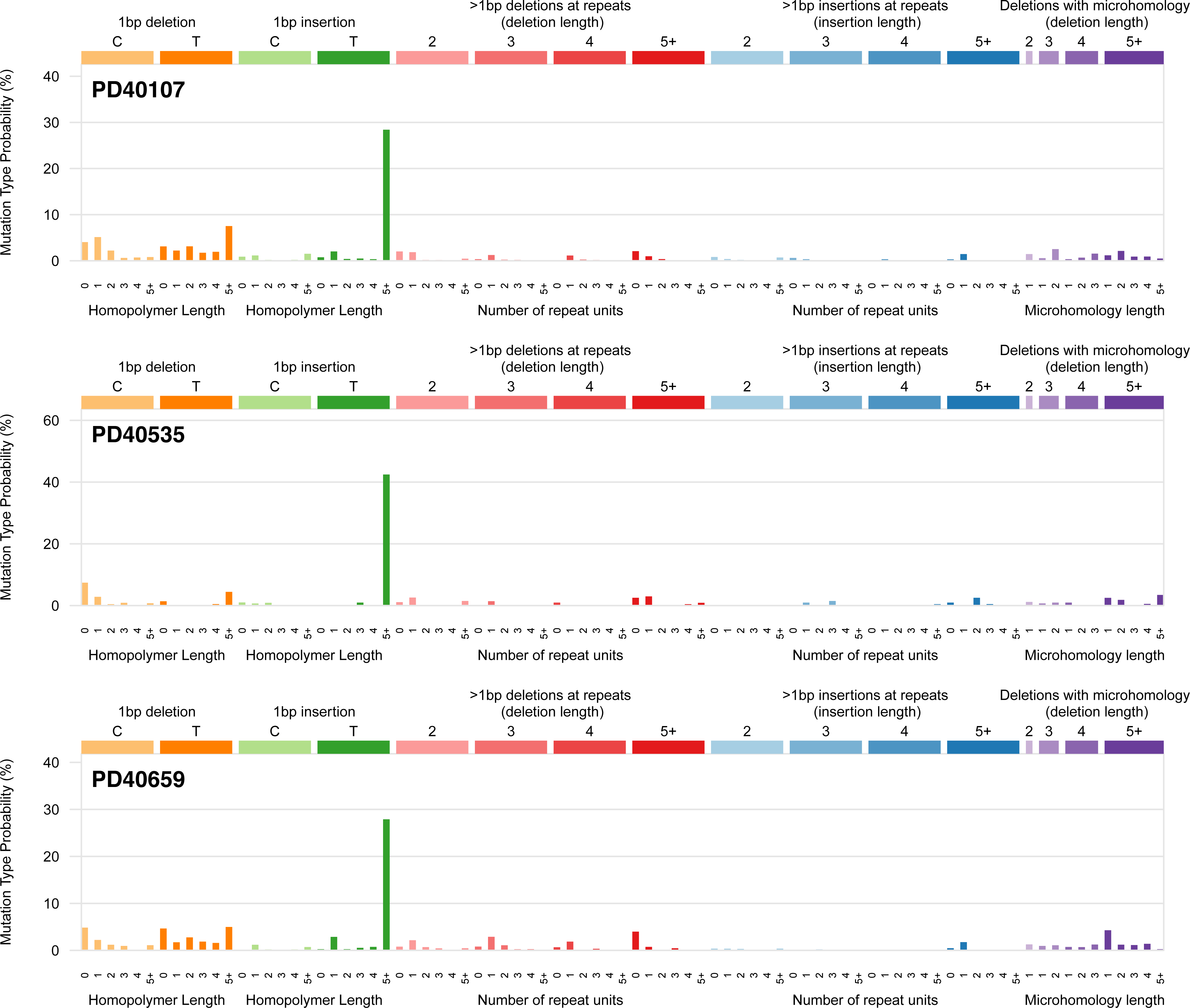

**Extended Data Figure 4.**
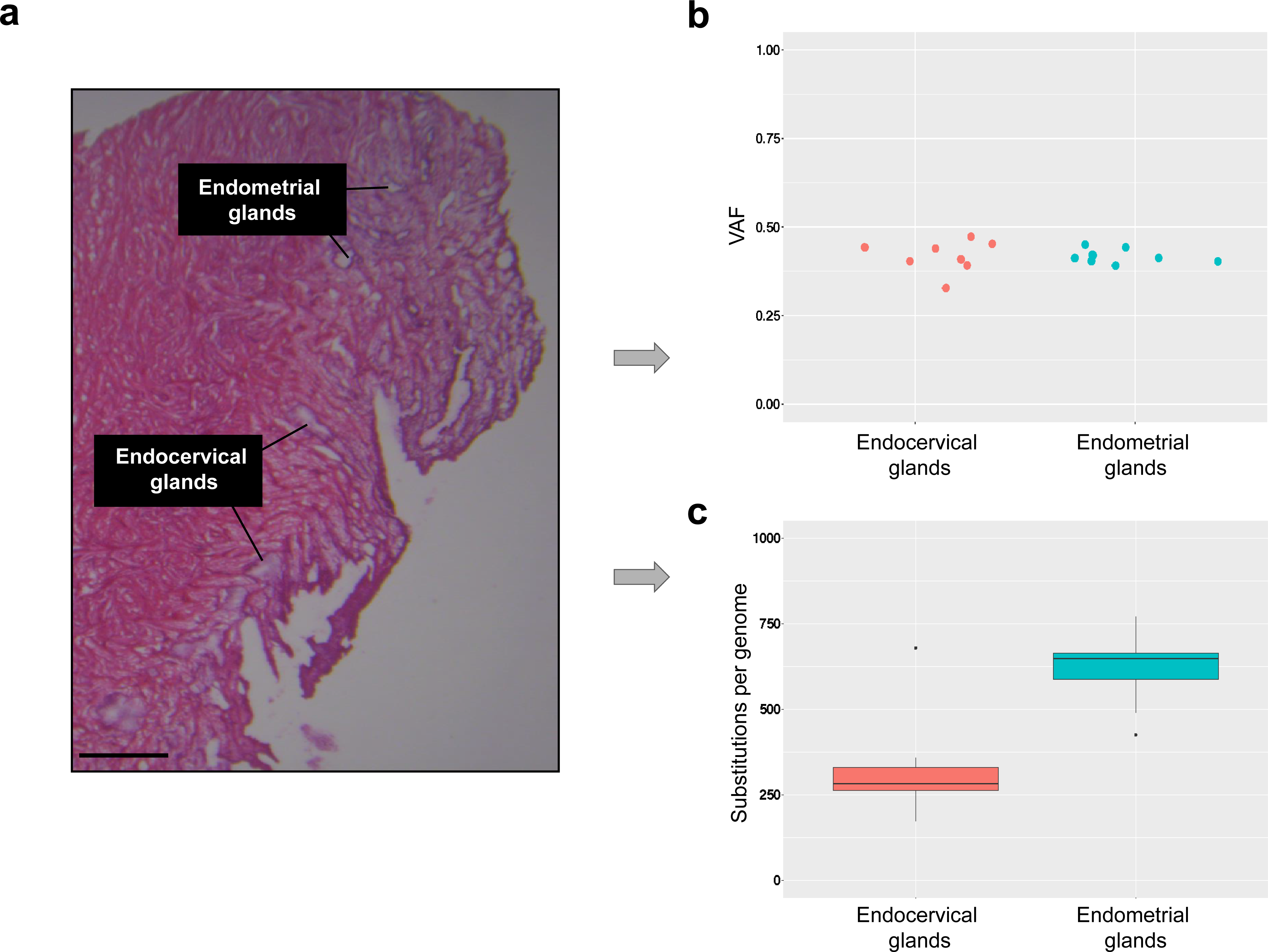
Comparison between normal endometrial and endocervical glands. (**a**) An overview histology image of an ∼2cm^3^ tissue biopsy sample from a 19-year-old donor (PD37506). The image shows normal endometrial and adjacent endocervical glands, which were subsequently micro-dissected. (**b**) Endometrial and endocervical glands with a similar median variant allele frequency (VAF) of substitutions were compared. (**c**) There was a ∼2-fold difference in the mutation burden between the two types of glands.

**Extended Data Figure 5.**
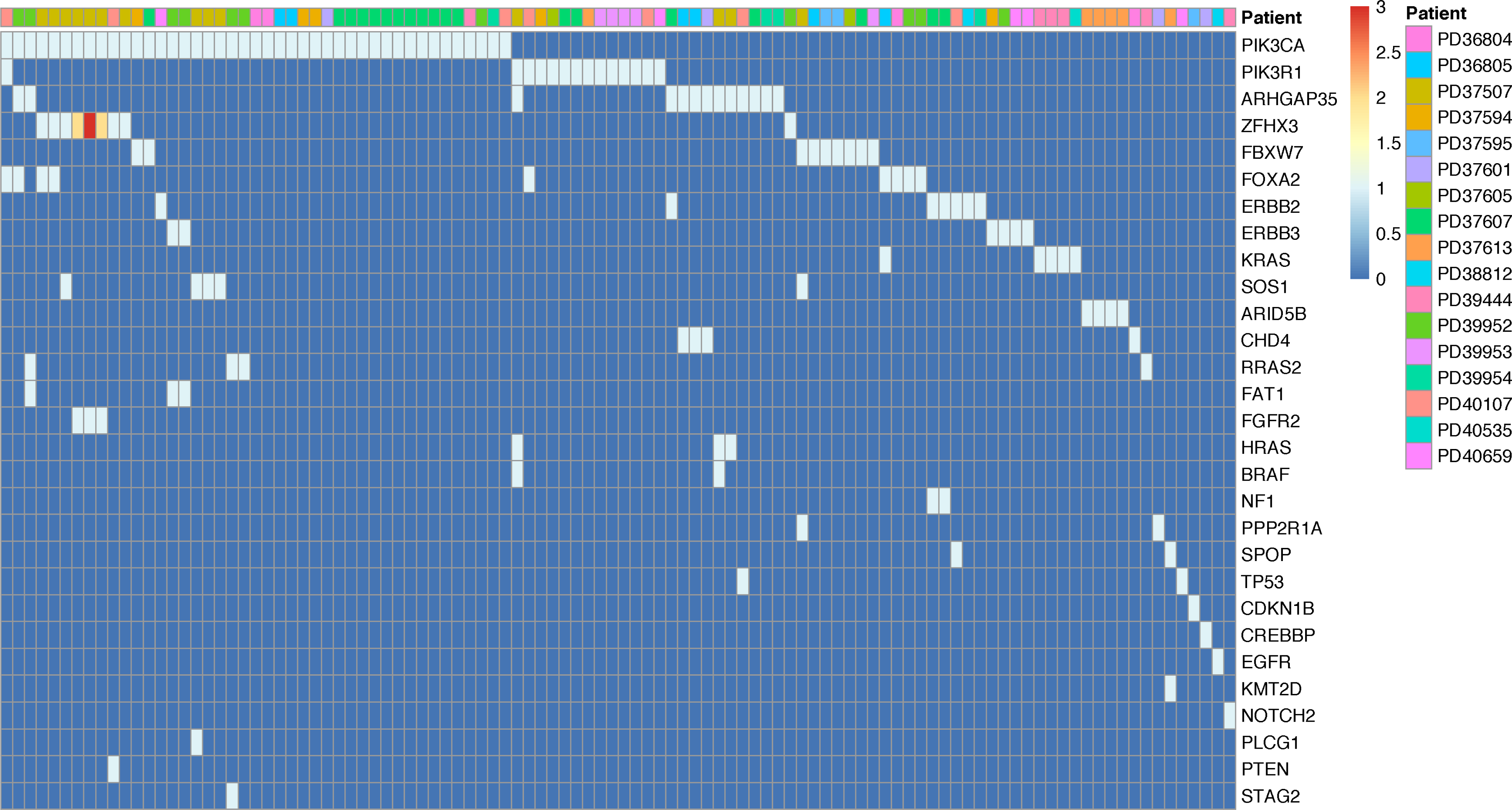
Oncoplot of all driver mutations and their distribution across individual endometrial gland samples and donors. Each cell represents an individual endometrial gland sample and is colour-coded to represent the total number of detected driver mutations (0-3). *PIK3CA* was the most frequently mutated gene with at least one mutation detected in 61% (11/18) of women. In some glands, these co-occurred with mutations in *ZFHX3*, *ARHGAP35*, *FGFR2*, *FOXA2* and other genes that are also selected for in endometrial cancer.

**Extended Data Figure 6.**
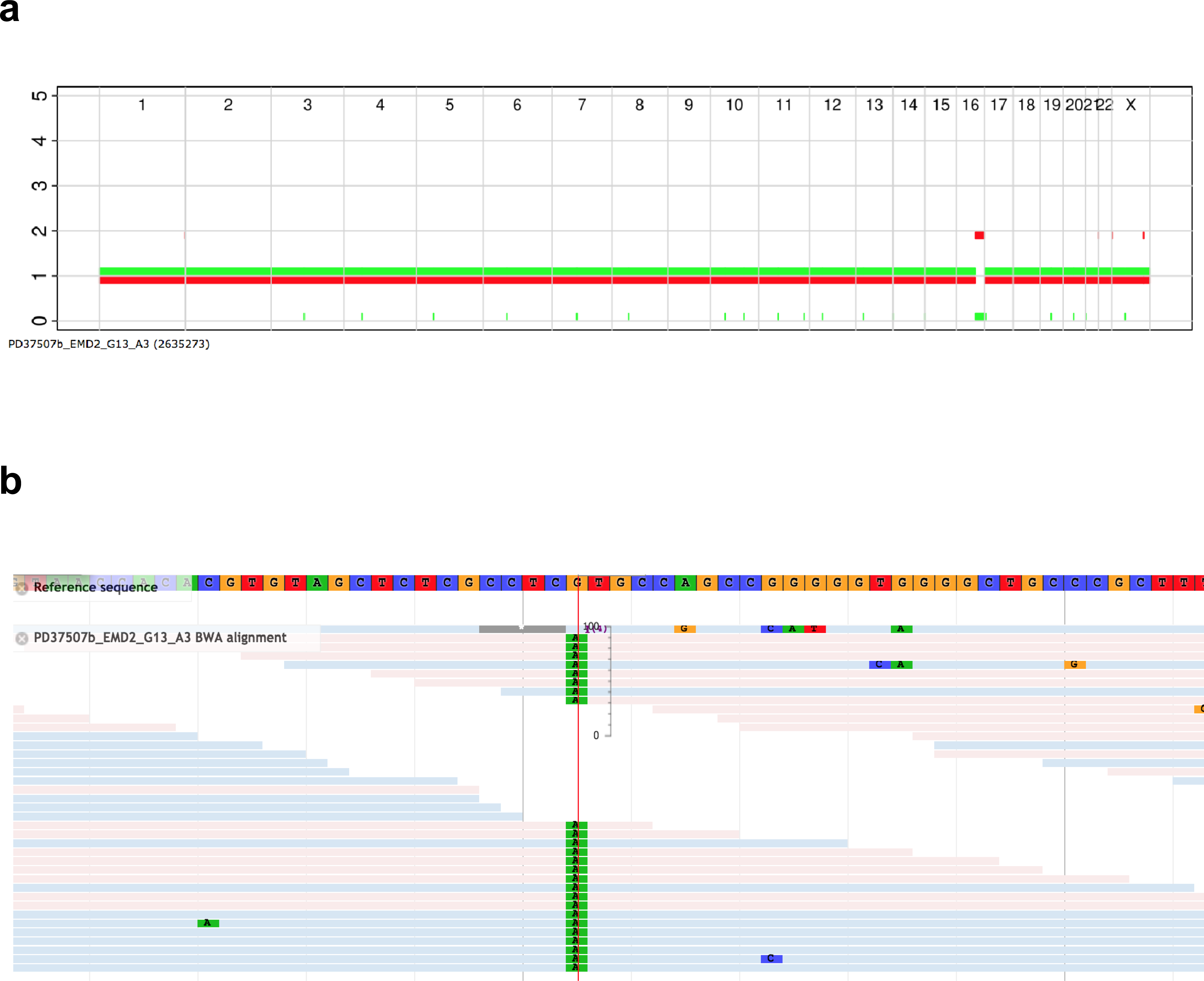
An example of copy-number neutral loss of heterozygosity (cnn-LOH) in a normal endometrial gland. (**a**) biallelic truncating mutation is seen in *ZFHX3* (p.R715*) with every read carrying the variant. (**b**) an associated cnn-LOH is observed on chromosome 16.

**Extended Data Figure 7.**
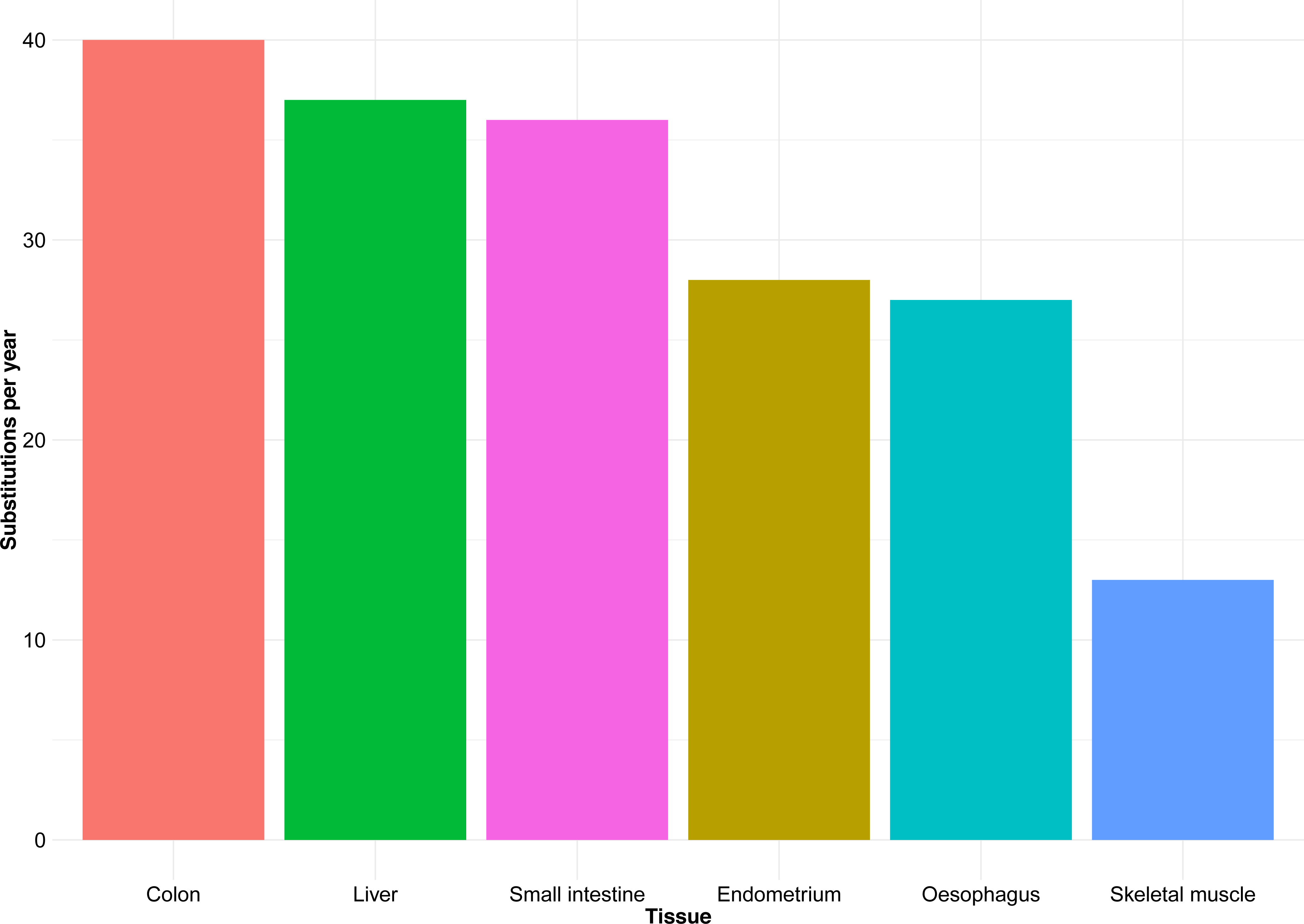
Comparison of mutation rates between endometrial epithelium and other cell types. The barplot shows a comparison of estimated mutilation rates (substitutions) for normal endometrial epithelial and other cell types from previously published studies (liver, colon and small intestine^4^, oesophagus^39^ and skeletal muscle^7^).

## SUPPLEMENTARY METHODS

### Sample collection

Anonymized snap-frozen endometrial tissue samples were obtained from five different cohorts.

Cohort 1: Samples from individuals PD37605, PD37601, PD37607, PD37613, PD37594 and PD37595 (age 29 to 46) were collected from women undergoing hysteroscopy examination as part of infertility assessment at the Tommy’s National Early Miscarriage Centre, University Hospitals Coventry and Warwickshire NHS Trust. Informed consent was obtained and biopsies collected and stored at the Arden Tissue Bank, University Hospitals Coventry and Warwickshire NHS Trust in line with the protocols approved by the NRES Committee South Central Southampton B (REC reference 12/SC/0526, 19/04/2013).

Cohort 2: Samples from individuals PD40535, PD39444, PD39953, PD39952, PD39954 and PD40107 (age 24 to 69) were collected from residual tissues from transplant organ donors with an informed consent obtained from donor’s family (REC reference: 15/EE/0152 NRES Committee East of England – Cambridge South).

Cohort 3: Individuals PD36804 and PD36805 (age 47 and 49), underwent total abdominal hysterectomy for benign non-endometrial pathologies and biopsies were collected, snap frozen and stored at the Human Research Tissue Bank, Cambridge University Hospitals NHS Foundation Trust in line with the protocols approved by the NRES Committee East of England (REC reference 11/EE/0011, 11/03/2011).

Cohorts 4 and 5: Samples from individuals PD37506, PD38812, PD37507 and PD40659 (age 19 to 81) were obtained at autopsy following death from non-gynaecological causes. The use of this material was approved by the London, Surrey Research Ethics Committee (REC reference 17/LO/1801, 26/10/2017) and East of Scotland Research Ethics Service (REC reference: 17/ES/0102, 27/07/2017).

All endometrial biopsies underwent formal pathology review, which confirmed benign histology.

### Laser-capture microdissection of endometrial glands

Frozen and paraffin sections were used for laser-capture microdissection (LCM). For frozen sections, endometrial tissue was embedded in optimal cutting temperature (OCT) compound. 14 to 20-micron thick sections were generated at -20°C to -23°C, mounted on to poly-ethylene naphtholate (PEN)-membrane slides (Leica), fixed with 70% ethanol, washed twice with phosphate-buffered saline (PBS), and stained with Gill’s haematoxylin and eosin for 20 and 10 seconds respectively.

For paraffin sections, frozen endometrial tissue was first thawed at 4°C for 10-15 minutes, then fixed in 70% ethanol and embedded in paraffin using standard histological tissue processing. 8 to 10-micron thick sections were subsequently cut, mounted on to PEN-membrane slides, and stained by sequential immersion in the following: xylene (two minutes, twice), ethanol (100%, 1 minute, twice), deionised water (1 minute, once), Gill’s haematoxylin (10-20 seconds), tap water (20 seconds, twice), eosin (10 seconds, once), tap water (10-20 seconds, once), ethanol (70%, 20 seconds, twice) and xylene or neo-clear xylene substitute (10-20 seconds, twice).

Using laser-capture microscope (Leica LMD7), individual endometrial glands were first visualised, then dissected (power 7, aperture 1, pulse 119 and speed 5) and collected into separate wells in a 96-well plate. Overview pre- and post-dissection images were taken. In addition, 200-500-µm^2^ sections of either myometrium, endometrial stroma or Fallopian tube epithelium were also obtained.

### Cell lysis, DNA extraction and whole genome sequencing of endometrial glands

20 μl of an in-house lysis buffer containing 30 mM Tris-HCl pH 8.0 (Sigma Aldrich), 0.5% Tween-20 (Sigma Aldrich), 0.5% NP-40/IGEPAL CA-630 (Sigma Aldrich) and 1.25 μg/ml Proteinase K (Qiagen) was added to each well, vortexed (30 seconds) and spun down at 18°C (one minute at 1500 rpm). Samples were subsequently incubated in a thermal cycler for 60 minutes at 50°C and 30 minutes at 75°C prior to storage at -80°C.

All samples in this study were processed using our recently developed low-input enzymatic fragmentation-based library preparation method^12^. Briefly, each 20 ul LCM lysate was mixed with 50 ul Ampure XP beads (Beckman Coulter) and 50 μl TE buffer (Ambion; 10 mM Tris-HCl, 1 mM EDTA) at room temperature. Following a 5 minute binding reaction and magnetic bead separation, genomic DNA was washed twice with 75% ethanol. Beads were resuspended in 26 μl TE buffer and the bead/genomic DNA slurry was processed immediately for DNA library construction. Each sample (26 μl) was mixed with 7 μl of 5X Ultra II FS buffer, 2 μl of Ultra II FS enzyme (New England BioLabs) and incubated on a thermal cycler for 12 minutes at 37°C then 30 minutes at 65°C. Following DNA fragmentation and A-tailing, each sample was incubated for 20 minutes at 20°C with a mixture of 30 μl ligation mix and 1 μl ligation enhancer (New England BioLabs), 0.9 μl nuclease-free water (Ambion) and 0.1 μl duplexed adapters (100 uM; 5’-ACACTCTTTCCCTACACGACGCTCTTCCGATC*T-3’, 5’-phos-GATCGGAAGAGCGGTTCAGCAGGAATGCCGAG-3’). Adapter-ligated libraries were purified using Ampure XP beads by addition of 65 μl Ampure XP solution (Beckman Coulter) and 65 μl TE buffer (Ambion). Following elution and bead separation, DNA libraries (21.5 μl) were amplified by PCR by addition of 25 μl KAPA HiFi HotStart ReadyMix (KAPA Biosystems), 1 μl PE1.0 primer (100 μM; 5’-AATGATACGGCGACCACCGAGATCTACACTCTTTCCCTACACGACGCTCTTCCGATC*T-3’) and 2.5 μl iPCR-Tag (40 μM; 5’-CAAGCAGAAGACGGCATACGAGATXGAGATCGGTCTCGGCATTCCTGCTGAACCGCTC TTCCGATC-3’) where ‘X’ represents one of 96 unique 8-base indexes The samples were then mixed and thermal cycled as follows: 98 °C for 5 minutes, then 12 cycles of 98 °C for 30 s, 65°C for 30 s, 72 °C for 1 minute and finally 72 °C for 5 minutes. Amplified libraries were purified using a 0.7:1 volumetric ratio of Ampure Beads (Beckman Coulter) to PCR product and eluted into 25 μl of nuclease-free water (Ambion). DNA libraries were adjusted to 2.4 nM and sequenced on the HiSeq X platform (illumina) according to the manufacturer’s instructions with the exception that we used iPCRtagseq (5’-AAGAGCGGTTCAGCAGGAATGCCGAGACCGATCTC-3’) to read the library index.

All LCM samples were subjected to whole genome sequencing of 15-40.3x, using 150 base pair clipped reads sequenced on HiSeq X platform (Illumina). For selected donors (PD36804, PD36805 and PD37506), bulk samples (fragments of uterus or cervix) were also whole genome sequenced.

### Variant calling

#### Substitutions

Sequencing data were aligned to the reference human genome (NCBI build 37) using Burrow-Wheeler Aligner (BWA-MEM)^41^. Duplicates were marked and removed and mapping quality thresholds were set at 30. Single base somatic substitutions were called using Cancer Variants through Expectation Maximization (CaVEMan) algorithm (major copy number 5, minor copy number 2)^42^. To exclude germline variants, matched normal samples (cervix, myometrium, Fallopian tube or endometrial stroma) were used to run the algorithm.

A set of previously described post-processing filters were subsequently applied:

1. to remove common single nucleotide polymorphisms, variants were filtered against a panel of 75 unmatched normal samples^42^;
2. to remove mapping artefacts associated with BWA-MEM, median alignment score of reads supporting a mutation should be greater than or equal to 140 (ASMD>=140) and fewer than half of the reads should be clipped (CLPM=0)^12^;
3. to remove artefacts that are specific to the library preparation for LCM samples, two additional filters were used. A fragment-based filter, which is designed to remove overlapping reads resulting from relatively shorter insert sizes allowed in this protocol that can lead to double counting of variants, and a cruciform filter, which removes erroneous variants that can be introduced due to the incorrect processing of cruciform DNA. For each variant, the standard deviation (SD) and median absolute deviation (MAD) of the variant position within the read was calculated separately for positive and negative strands reads. If a variant was supported by a low number of reads for one strand, the filtering was based on the statistics calculated from the reads derived from the other strand and it was required that either: (a) ≤ 90% of supporting reads report the variant within the first 15% of the read as determined from the alignment start, or (b) that the MASD >0 and SD>4. Where both strands were supported by sufficient reads, it was required for both strands separately to either: (a) ≤90% of supporting reads report the variant within the first 15% of the read as determined from the alignment start, (b) that the MAD>2 and SD>2, or (c) that at least one strand has fulfilled the criteria MAD>1 and SD>10.

#### Validation experiments and sensitivity

To validate somatic variants, for selected donors, pairs of biological ‘near-replicates’ were obtained. For these experiments, we collected two samples from the same endometrial gland which was identified on two or more consecutive levels using z-stacking approach; each sample was processed separately with an independent DNA extraction, library preparation and whole genome sequencing. As these samples were obtained from the same glands, they would represent derivatives of the same clone and therefore the same sensitivity would be assumed in both samples in each pair. The maximum likelihood estimate for sensitivity (*s*) was then calculated as follows:

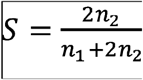

where *n*_1_ is the number of variants called only in one of the two LCM samples and *n*_2_ is the number of variants called in both LCM samples in each pair. Using this approach, the mean sensitivity of somatic mutation variant calling was estimated at >86% (range 0.70-0.95%).

#### Indels

Insertions and deletions were called using cgpPindel^43,44^. To remove germline variants the algorithm was run with the same matched normal samples that were used for calling substitutions. Post-processing filters were applied as previously described^42^. In addition, a ‘Qual’ filter (the sum of the mapping qualities of the supporting reads) of at least 300 and a depth cut-off of at least 15 reads were used.

#### Copy number variants and structural variants

Allele-specific copy number profiles were reconstructed for the endometrial gland samples by ASCAT^45,46^ using matched samples as described above, ploidy of 2 and contamination with other cell types of 10%. Only samples with a minimum coverage of 15X and above were used. All putative copy number changes were visually inspected for copy number profiles on Jbrowse^47^.

Structural variants (SVs) in endometrial glands were called using matched samples (as described above) with the Breakpoints Via Assembly (BRASS) algorithm and further annotated by GRASS (https://github.com/cancerit/BRASS). Potential SVs are detected for the sample of interest and read-pairs clusters supporting the SV are used for breakpoint sequence de novo assembly. Absence of supporting evidence in the matched control indicates that the SV was acquired in the sample of interest. The isolation of minute amounts of DNA for sequencing in combination with the LCM enzymatic fragmentation-based library preparation procedure introduces additional artefacts and additonal post-processing filtering was performed in two phase:

#### Further annotation of SVs with statistics that detect LCM specific artefacts

All SVs detected by BRASS were further annotated by AnnotateBRASS. Each SV is defined by two breakpoints and their genomic coordinates.

A. The following statistics were determined for each breakpoint separately:
  1. The total number of reads supporting the SV.
  2. The total number of unique reads supporting the SV, based on alignment position and read orientation.
  3. The standard deviation of the alignment positions of reads supporting the SV.
  4. The number of chromosomes, based on read-pairs not supporting the SV, to which one read mapped while the mate-read aligned to the SV breakpoint.
  5. The number of reads supporting the SV that had an alternative alignment (XA-tag).
  6. The number of reads supporting the SV that had an alternative alignment score (XS-tag) similar to the current alignment score.
  7. The percentage of read-pairs not supporting the SV with a discordant inferred insert size (default: ≥ 1000bp).
B. A wider search for read-pairs supporting the SV is intiated and the following statistics were calculated for each breakpoint separately:
  1. The total number of reads supporting the SV.
  2. The total number of unique reads supporting the SV, based on alignment position and read orientation.
  3. The standard deviation of the alignment positions of reads supporting the SV.
  4. The number of reads supporting the SV that had an alternative alignment.
  5. The number of reads supporting the SV that had and alternative alignment score similar to the current alignment score.
C. Reads spanning the SV breakpoints are often clipped. Clipped sequences of sufficient length can be aligned to other positions on the genome (i.e., supplementary alignment) and it is expected that these align to the proximity of the other SV breakpoint. Based on the clipping positions and supplementary alignments the following was determined for each SV:
  1. Whether the clipped sequences of read-pairs spanning a SV breakpoint align in the proximity of the other SV breakpoint.
  2. Whether the clipping within read-pairs supporting the SV occurred at roughly the same genomic position (default: all clipping positions occurred within 10bp of each other).
D. BRASS uses a single matched control and a panel of normals (PoN, bulk WGS) to determine whether a SV is somatic. SVs observed in the sample of interest but not in the matched control or PoN are considered somatic. However, due to the difference in library preparation and the variance of spatial genomic coverage observed it is not always possible to accurately assess the validity of the SV. Two different approaches were implemented to determine whether the SV is somatic:
  1. A wider search in the matched control sample was performed to search for read-pairs that could support the SV. The SV was still considered detected in case the discovered read-pairs were insufficient for breakpoint sequence de novo assembly.
  2. Additional controls can be defined in case multiple samples have been isolated for the same individual. Samples from the same individual with little genetic relationship, as determined from the SNVs and indels, can be used as controls to determine whether te detected SV is germline or a recurrent artifact.

#### Post-hoc filtering of SVs based on a combination of the above statistics

SVs were further filtered based on the described statistics. The optimal set of statistics and their most practical thresholds depends on the achieved coverage and stringency of filtering desired. At default the following criteria were used for detecting somatic SVs:

1. For each breakpoint there must be ≥ 4 unique reads supporting the SV (**A.2**).
2. The alignment position standard deviation must be > 0 (**A.3**).
3. At each breakpoint there are read-pairs not supporting the SV that map to < 5 other chromosomes (**A.4**).
4. The total number of chromosomes mapped to by read-pairs not supporting the SV for both breakpoints should be < 7 (**A.4**).
5. The percentage of reads supporting the SV with alternative alignments or alternative alignments with similar alignment scores should be ≤ 50% for both SV breakpoints separately (**A.5-A.6**).
6. The percentage of discordant read-pairs not supporting the SV should be ≤ 7.5% of total read-pairs for both SV breakpoints separately (**A.7**).
7. For the wider search of SV-supporting read-pairs the same thresholds apply as under criteria 1-6 (**B.1-B.5**).
8. There are no read-pairs in the matched control that support the SV (**C.1**).
9. The SV is not detected in any of the other control samples, or there were ≤ 2 samples carrying the same SV and the proportion of control samples carrying the SV was < 1/3 of the defined control set (**C.2**).
10. It was not allowed for read-pairs supporting the SV to have widely divergent clipping positions in terms of genomic location for both SV breakpoints separately (**D.2**).

#### Detection of driver mutations

Analysis of driver variants in the normal endometrial glands was performed in two parts. First, filtered CaVEMan and Pindel variants were intersected against a previously published list of 369 genes that are under selection in human cancers^30^. All non-synonymous mutations were annotated to indicate mode of action using a Cancer Gene Census (719 genes) and a catalogue of 764 genes (https://www.cancergenomeinterpreter.org). Truncating variants (nonsense, frameshift and essential splice), which resided in recessive/tumour-suppressor genes (TSG) were declared likely drivers. Missense mutations in recessive/TSG and dominant/oncogenes were triaged against a database of validated hotspot mutations (http://www.cbioportal.org/mutation_mapper). All mutations that were shown to be known mutational hotspots or ‘likely oncogenic’ were declared drivers. In addition, identified activating mutations in mutational hotspots in genes *RRAS2* and *SOS1*, involving the RAS/MAPK pathway were declared as likely drivers.

Second, to identify genes that are under positive selection in normal endometrium we used the dN/dS^30^ method that is based on the observed:expected ratios of non-synonymous:synonymous mutations. The analysis was carried out for the whole genome (q<0.05 and q<0.01) and for 369 known cancer genes^30^ (RHT, restricted hypothesis testing, q<0.05). Eleven genes were found to be under positive selection in normal endometrial glands. The output of this analysis was also used to assess whether missense mutations in genes that are under positive selection in normal and/or malignant endometrium (*PIK3CA, ERBB2, ERBB3, FBXW7* and *CHD4*) but are not known mutational hotspots, are likely to be drivers. If q-value was <0.05, we used the following calculation to assess the likelihood of a variant being a driver:

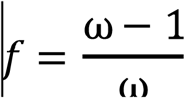

If *f* was ≥ 0.95, then all missense mutations in that gene were declared likely drivers.

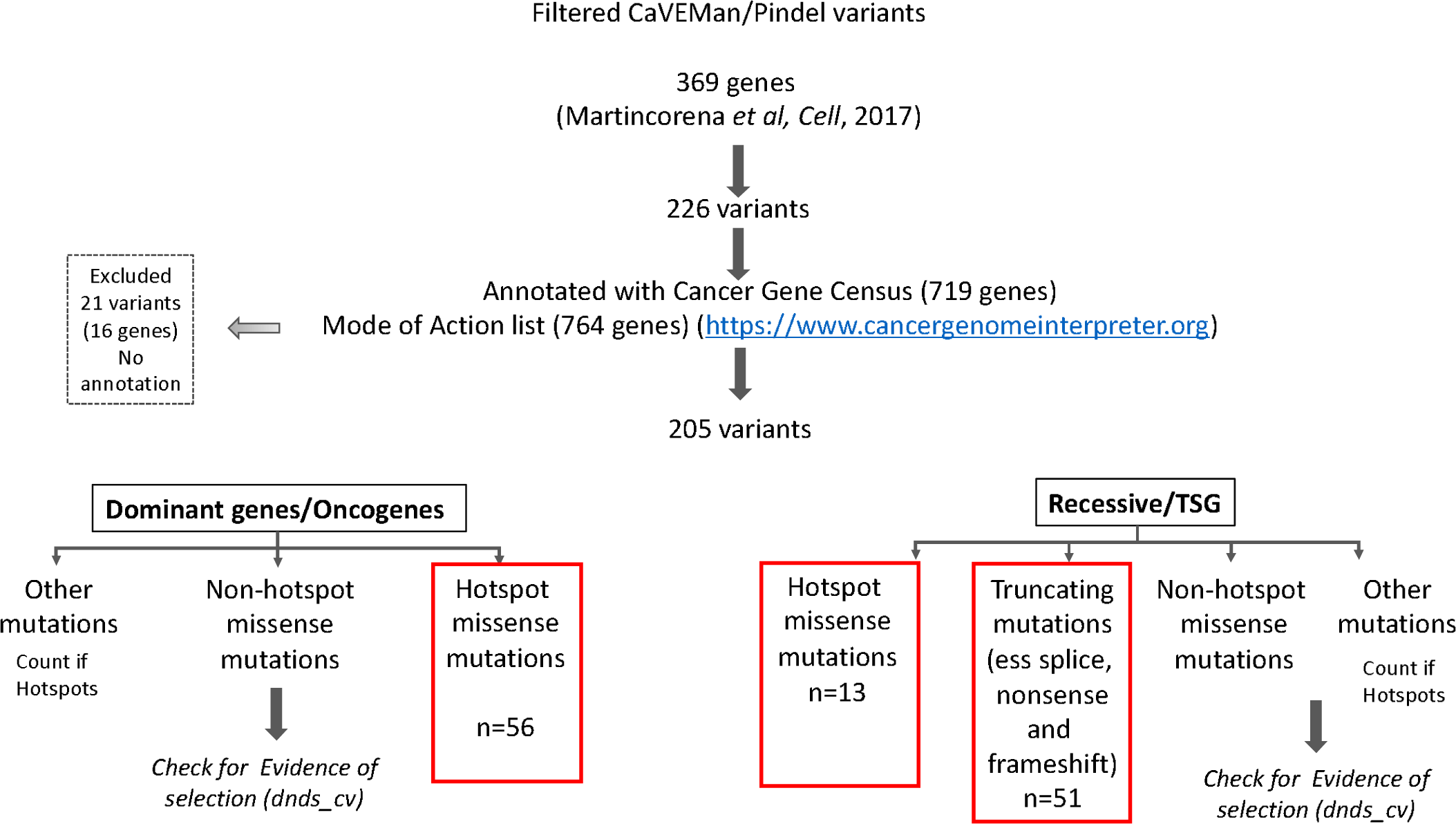

To compare patterns of selection in normal endometrial epithelium and cancer, we performed dNdS analysis on publicly available calls from the The Cancer Genome Atlas, TCGA.

#### Phylogenetic tree reconstruction

Phylogenies for endometrial glands were reconstructed for seventeen donors. Due to the low number of available samples, donor PD38812 was not included in this analysis. We first generated trees using substitutions called by CaVEMan; matched normal samples were used to exclude germline variants and post-processing filters were applied as above. Final variants were recalled in all samples from each donor using an in-house re-genotyping algorithm (cgpVAF). Variants with a VAF>0.3 were noted to be present (‘1’), VAF<0.1 absent (‘0’) and between 0.1 and 0.3 as ambiguous (‘?’). This approach excludes private sub-clonal variants from the tree building. The tree was reconstructed using a maximum parsimony approach^48^ and branch support was calculated using 1000 bootstrap replicates. Nodes with a confidence lower than 50 were collapsed into polytomies and branch lengths of the collapsed tree were determined by the number of assigned substitutions.

The constructed phylogenies were validated using indels called by Pindel and filtered as above. The same approach was applied for the final indel matrices. Although the lower number of indels resulted in more polytomous tree, the overall tree topologies were reconcilable with those generated using substitutions.

Cancer driver mutations, copy number and structural variants were annotated manually in the trees.

#### Mutational signature analysis

Mutational signature extraction was performed using mutations assigned to every branch of the reconstructed phylogenetic trees and each branch was treated as an individual sample. Such approach allows characterisation and differentiation of specific mutational processes that were operative at various times in individual glands. Substitutions were first categorised into 96 classes following the method used by the Mutational Signature working group of the Pan Cancer Analysis of Whole Genomes (PCAWG)^2^. Single base substitution (SBS) signatures were then extracted using the HDP package (https://github.com/nicolaroberts/hdp) that utilises hierarchical Bayesian Dirichlet process. Code and the input mutations are available at https://github.com/LuizaMoore/Endometrium. SBS signature analysis was performed in 3 steps: extraction, deconvolution and re-attribution. First, extraction was performed with conditioning on the set of mutational signatures that have been previously reported in endometrial cancer by PCAWG: SBS1, SBS2, SBS3, SBS5, SBS6, SBS7a, SBS7b, SBS10a, SBS10b, SBS13, SBS14, SBS15, SBS20, SBS21, SBS26, SBS28, SBS30, SBS44 and SBS54 (reference). Such an approach not only allows discovery of new mutational signatures, but also simultaneous matching to the provided reference signatures. The extraction was run with 50,000 burn-in iterations (parameter ‘burnin’), with a spacing of 500 iterations (parameter ‘space’) and 250 samples/iterations were collected (parameter ‘n’). After each Gibbs sampling iteration, 3 iterations of concentration parameter sampling were performed (parameter ‘cpiter’) and components were extracted (Cos merge 0.9, sample 2).

Two reference signatures were extracted: SBS1 (P1, cosine similarity 0.997) and SBS5 (P5, cosine similarity 0.983), which were added to the ‘deconvolution’ catalogue. Other extracted components (New 1 = N1, New 2 = N2, New 3 = N3 and Component 0 = Comp 0) that did not fit the provided set of 19 reference signatures were examined for similarity to the full set of 60 reference SBS signatures (PCAWG)^2^. Component N2 showed high cosine similarity to SBS18 (0.968), therefore did not require further deconvolution and was added to the ‘deconvolution’ catalogue. No other HDP components showed cosine similarity of >0.95 to any of the reference signatures and therefore required further deconvolution. The final ‘deconvolution’ catalogue comprising the three extracted SBS signatures (SBS1, SBS5 and SBS18) was then used to decipher all other components. Final SBS signature exposures were re-calculated and signatures re-attributed to individual samples.

Indels were classified using PCAWG method^2^ and composite mutational spectra were generated for each donor. However, given the relatively low numbers of indels, no formal signature extraction was performed.

### Calculations of mutation burden and estimation of mutation rate

To account for the non-independent sampling per patient we used mixed effects models. We tested features with a known effect on mutation burden or endometrial cancer risk; age, Read depth & VAF, BMI and Parity. All statistical analyses were performed in R and are summarised in the Supplementary Results.

### Data availability

Whole genome sequencing data are deposited in the European Genome 715 Phenome Archive (EGA) with accession number 716 EGAS00001002471.

### Code availability

Code for statistical analyses on total substitution and driver mutation burdens is included in the supplementary material. Code for mutational signature extraction is deposited on GitHub at https://github.com/LuizaMoore/Endometrium. All other code is available on request.

## REFERENCES

1 Alexandrov, L. B. Signatures of mutational processes in human cancer. Nature 500, 415–421, doi:10.1038/nature12477 (2013).

2 Alexandrov, L. The Repertoire of Mutational Signatures in Human Cancer. Environ Mol Mutagen 59, 25–25 (2018).

3 Stratton, M. R., Campbell, P. J. & Futreal, P. A. The cancer genome. Nature 458, 719-724, doi:10.1038/nature07943 (2009).

4 Blokzijl, F. et al. Tissue-specific mutation accumulation in human adult stem cells during life. Nature 538, 260-264, doi:10.1038/nature19768 (2016).

5 Lee-Six, H. et al. Population dynamics of normal human blood inferred from somatic mutations. Nature 561, 473-478, doi:10.1038/s41586-018-0497-0 (2018).

6 Bae, T. et al. Different mutational rates and mechanisms in human cells at pregastrulation and neurogenesis. Science 359, 550-+, doi:10.1126/science.aan8690 (2018).

7 Franco, I. et al. Somatic mutagenesis in satellite cells associates with human skeletal muscle aging. Nat Commun 9, 800, doi:10.1038/s41467-018-03244-6 (2018).

8 Osorio, F. G. et al. Somatic Mutations Reveal Lineage Relationships and Age-Related Mutagenesis in Human Hematopoiesis. Cell Rep 25, 2308-2316 e2304, doi:10.1016/j.celrep.2018.11.014 (2018).

9 Martincorena, I. et al. Tumor evolution. High burden and pervasive positive selection of somatic mutations in normal human skin. Science 348, 880-886, doi:10.1126/science.aaa6806 (2015).

10 Martincorena, I. Somatic mutant clones colonize the human esophagus with age. Science 362, 911-917, doi:10.1126/science.aau3879 (2018).

11 Suda, K. et al. Clonal Expansion and Diversification of Cancer-Associated Mutations in Endometriosis and Normal Endometrium. Cell Rep 24, 1777-1789, doi:10.1016/j.celrep.2018.07.037 (2018).

12 Lee-Six, H., et al. The landscape of somatic mutation in normal colorectal epithelial cells., <bioRxiv 416800> (2018).

13 Schmitt, M. W. et al. Detection of ultra-rare mutations by next-generation sequencing. P Natl Acad Sci USA 109, 14508-14513, doi:10.1073/pnas.1208715109 (2012).

14 Hoang, M. L. et al. Genome-wide quantification of rare somatic mutations in normal human tissues using massively parallel sequencing. Proc Natl Acad Sci U S A 113, 9846-9851, doi:10.1073/pnas.1607794113 (2016).

15 Michael A. Lodato, R. E. R., Craig L. Bohrson, Michael E. Coulter. Aging and neurodegeneration are associated with increased mutations in single human neurons. Science 359, 555-559, doi:10.1126/science.aao4426 (2018).

16 Lodato, M. A. et al. Somatic mutation in single human neurons tracks developmental and transcriptional history. Science 350, 94-98, doi:10.1126/science.aab1785 (2015).

17 Jaiswal, S. et al. Age-related clonal hematopoiesis associated with adverse outcomes. N Engl J Med 371, 2488-2498, doi:10.1056/NEJMoa1408617 (2014).

18 Genovese, G. et al. Clonal hematopoiesis and blood-cancer risk inferred from blood DNA sequence. N Engl J Med 371, 2477-2487, doi:10.1056/NEJMoa1409405 (2014).

19 Xie, M. et al. Age-related mutations associated with clonal hematopoietic expansion and malignancies. Nat Med 20, 1472-1478, doi:10.1038/nm.3733 (2014).

20 McKerrell, T. et al. Leukemia-associated somatic mutations drive distinct patterns of age-related clonal hemopoiesis. Cell Rep 10, 1239-1245, doi:10.1016/j.celrep.2015.02.005 (2015).

21 Zink, F. et al. Clonal hematopoiesis, with and without candidate driver mutations, is common in the elderly. Blood 130, 742-752, doi:10.1182/blood-2017-02-769869 (2017).

22 Cousins, F. L., O, D. F. & Gargett, C. E. Endometrial stem/progenitor cells and their role in the pathogenesis of endometriosis. Best Practice & Research Clinical Obstetrics & Gynaecology 50, 27-38, doi:10.1016/j.bpobgyn.2018.01.011 (2018).

23 Gargett, C. E., Schwab, K. E. & Deane, J. A. Endometrial stem/progenitor cells: the first 10 years. Hum Reprod Update 22, 137-163, doi:10.1093/humupd/dmv051 (2016).

24 Kaitu’u-Lino, T. J., Ye, L. & Gargett, C. E. Reepithelialization of the uterine surface arises from endometrial glands: evidence from a functional mouse model of breakdown and repair. Endocrinology 151, 3386-3395, doi:10.1210/en.2009-1334 (2010).

25 Tempest, N., Maclean, A. & Hapangama, D. K. Endometrial Stem Cell Markers: Current Concepts and Unresolved Questions. Int J Mol Sci 19, doi:10.3390/ijms19103240 (2018).

26 Le Gallo, M. & Bell, D. W. The emerging genomic landscape of endometrial cancer. Clin Chem 60, 98-110, doi:10.1373/clinchem.2013.205740 (2014).

27 Morice, P., Leary, A., Creutzberg, C., Abu-Rustum, N. & Darai, E. Endometrial cancer. Lancet 387, 1094-1108, doi:10.1016/S0140-6736(15)00130-0 (2016).

28 Onstad, M. A., Schmandt, R. E. & Lu, K. H. Addressing the Role of Obesity in Endometrial Cancer Risk, Prevention, and Treatment. J Clin Oncol 34, 4225-4230, doi:10.1200/JCO.2016.69.4638 (2016).

29 Setiawan, V. W. et al. Type I and II Endometrial Cancers: Have They Different Risk Factors? Journal of Clinical Oncology 31, 2607-+, doi:10.1200/Jco.2012.48.2596 (2013).

30 Martincorena, I. et al. Universal Patterns of Selection in Cancer and Somatic Tissues. Cell 171, 1029-1041 e1021, doi:10.1016/j.cell.2017.09.042 (2017).

31 Getz, G. Integrated genomic characterization of endometrial carcinoma (vol 497, pg 67, 2013). Nature 500, doi:10.1038/nature12325 (2013).

32 Anglesio, M. S. et al. Cancer-Associated Mutations in Endometriosis without Cancer. N Engl J Med 376, 1835-1848, doi:10.1056/NEJMoa1614814 (2017).

33 Lac, V. et al. Iatrogenic endometriosis harbors somatic cancer-driver mutations. Hum Reprod, doi:10.1093/humrep/dey332 (2018).

34 Lopez-Garcia, C., Klein, A. M., Simons, B. D. & Winton, D. J. Intestinal stem cell replacement follows a pattern of neutral drift. Science 330, 822-825, doi:10.1126/science.1196236 (2010).

35 Snippert, H. J. et al. Intestinal crypt homeostasis results from neutral competition between symmetrically dividing Lgr5 stem cells. Cell 143, 134-144, doi:10.1016/j.cell.2010.09.016 (2010).

36 Rouhani, F. J. et al. Mutational History of a Human Cell Lineage from Somatic to Induced Pluripotent Stem Cells. PLoS Genet 12, e1005932, doi:10.1371/journal.pgen.1005932 (2016).

37 Alexandrov, L. B. et al. Clock-like mutational processes in human somatic cells. Nat Genet 47, 1402-1407, doi:10.1038/ng.3441 (2015).

38 Zhang, Y. Y. L., Kucherlapati, M. A Pan-Cancer Compendium of Genes Deregulated by Somatic Genomic Rearrangement across More Than 1,400 Cases. Cell Rep. 24, 515–527, doi:10.1016/j.celrep.2018.06.025 (2018).

39 Martincorena, I. et al. Somatic mutant clones colonize the human esophagus with age. Science 362, 911-917, doi:10.1126/science.aau3879 (2018).

40 CRUK https://www.cancerresearchuk.org/health-professional/cancer-statistics/statistics-by-cancer-type/uterine-cancer/risk-factors#heading-Zero Accessed on 19/12/2018

## REFERENCES FOR SUPPLEMENTARY METHODS

41 Li, H. & Durbin, R. Fast and accurate short read alignment with Burrows-Wheeler transform. Bioinformatics 25, 1754-1760, doi:10.1093/bioinformatics/btp324 (2009).

42 Nik-Zainal, S. et al. Mutational processes molding the genomes of 21 breast cancers. Cell 149, 979-993, doi:10.1016/j.cell.2012.04.024 (2012).

43 Raine, K. M. et al. cgpPindel: Identifying Somatically Acquired Insertion and Deletion Events from Paired End Sequencing. Curr Protoc Bioinformatics 52, 15 17 11-12, doi:10.1002/0471250953.bi1507s52 (2015).

44 Ye, K., Schulz, M. H., Long, Q., Apweiler, R. & Ning, Z. Pindel: a pattern growth approach to detect break points of large deletions and medium sized insertions from paired-end short reads. Bioinformatics 25, 2865-2871, doi:10.1093/bioinformatics/btp394 (2009).

45 Van Loo, P. et al. Allele-specific copy number analysis of tumors. Proc Natl Acad Sci U S A 107, 16910-16915, doi:10.1073/pnas.1009843107 (2010).

46 Raine, K. M. et al. ascatNgs: Identifying Somatically Acquired Copy-Number Alterations from Whole-Genome Sequencing Data. Curr Protoc Bioinformatics 56, 15 19 11-15 19 17, doi:10.1002/cpbi.17 (2016).

47 Buels, R. et al. JBrowse: a dynamic web platform for genome visualization and analysis. Genome Biol 17, 66, doi:10.1186/s13059-016-0924-1 (2016).

48 Hoang, D. T. et al. MPBoot: fast phylogenetic maximum parsimony tree inference and bootstrap approximation. BMC Evol Biol 18, 11, doi:10.1186/s12862-018-1131-3 (2018).

